# Interactions between influenza A virus nucleoprotein and gene segment UTRs facilitate selective modulation of viral gene expression

**DOI:** 10.1101/2022.01.09.475567

**Authors:** Meghan Diefenbacher, Timothy JC Tan, David LV Bauer, Beth Stadtmueller, Nicholas C. Wu, Christopher B. Brooke

**Affiliations:** Department of Microbiology, University of Illinois at Urbana-Champaign, Urbana, Illinois, United States of America; Center for Biophysics and Quantitative Biology, University of Illinois at Urbana-Champaign, Urbana, Illinois, United States of America; RNA Virus Replication Laboratory, The Francis Crick Institute, London, United Kingdom; Department of Biochemistry, University of Illinois at Urbana-Champaign, Urbana, Illinois, United States of America; Department of Biomedical and Translational Sciences, Carle Illinois College of Medicine, University of Illinois at Urbana-Champaign, Urbana, Illinois, United States of America; Carl R. Woese Institute for Genomic Biology, University of Illinois at Urbana-Champaign, Urbana, Illinois, United States of America

## Abstract

The influenza A virus (IAV) genome is divided into eight negative-sense, single-stranded RNA segments. Each segment exhibits a unique level and temporal pattern of expression, however the exact mechanisms underlying the patterns of individual gene segment expression are poorly understood. We previously demonstrated that a single substitution in the viral nucleoprotein (NP:F346S) selectively modulates neuraminidase (NA) gene segment expression while leaving other segments largely unaffected. Given what is currently known about NP function, there is no obvious explanation for how changes in NP can selectively modulate the replication of individual gene segments. We found that the specificity of this effect for the NA segment is virus strain specific and depends on the UTR sequences of the NA segment. While the NP:F346S substitution did not significantly alter the RNA binding or oligomerization activities of NP *in vitro*, it specifically decreased the ability of NP to promote NA segment vRNA synthesis. In addition to NP residue F346, we identified two other adjacent aromatic residues in NP (Y385 & F479) capable of similarly regulating NA gene segment expression, suggesting a larger role for this domain in gene-segment specific regulation. Our findings reveal a new role for NP in selective regulation of viral gene segment replication and demonstrate how the expression patterns of individual viral gene segments can be modulated during adaptation to new host environments.

**Author summary:** Influenza A virus (IAV) is a respiratory pathogen that remains a significant source of morbidity and mortality. Escape from host immunity or emergence into new host species often requires mutations that modulate the functional activities of the viral glycoproteins hemagglutinin (HA) and neuraminidase (NA) which are responsible for virus attachment to and release from host cells, respectively. Maintaining the functional balance between the activities of HA and NA is required for fitness across multiple host systems. Thus, selective modulation of viral gene expression patterns may be a key determinant of viral immune escape and cross-species transmission potential. We identified a novel mechanism by which the viral nucleoprotein (NP) gene can selectively modulate NA segment replication and gene expression through interactions with the segment UTR. Our work highlights an unexpected role for NP in selective regulation of expression from the individual IAV gene segments.

## Introduction

Influenza A virus (IAV) is a major respiratory pathogen that causes seasonal epidemics and occasional pandemics that result in substantial morbidity and mortality (1). The genome of IAV is divided into eight negative sense, single-stranded RNA segments that encode one or more viral proteins (2). These negative-sense genomic RNAs (vRNAs) are used as templates to synthesize both the mRNA needed for protein synthesis and the positive-sense replicative intermediates (cRNAs) for genome replication (2). The individual IAV gene segments vary in both overall expression levels and timing, but the specific factors that govern this variation are poorly understood (3, 4).

Each IAV gene segment consists of one or more open reading frames (ORFs) flanked by untranslated regions (UTRs) (2). The UTRs consist of both segment specific sequences, and highly conserved sequences at the 3’ and 5’ termini which interact with one another to form the viral promoter (2). Previous studies have established roles for segment-specific sequences within the UTRs in modulating gene expression in a segment-specific manner (5–10). In virions and within infected cells, the gene segments are maintained as viral ribonucleoprotein complexes (vRNPs) in which the viral RNA is bound along its length by nucleoprotein (NP) and is associated with the viral RNA dependent RNA polymerase (RdRp) (2).

NP is a highly conserved (11) and multi-functional protein. To perform its integral role in vRNP formation, NP has two major known activities: RNA binding and oligomerization. NP binds RNA non-specifically through a positively charged groove located between its head and body domains (12, 13). Oligomerization of individual NP protomers occurs through the insertion of a C-terminal tail loop into the receptor groove of the neighboring protomer (12,14,15). As a key component of the vRNP complex, NP plays an essential role in vRNA replication and mRNA transcription. NP is hypothesized to act as an elongation factor for the viral polymerase as only short transcripts (<100nts) can be generated in its absence or in the presence of binding/oligomerization-deficient NP mutants (15). NP facilitates the import and export of vRNPs from the nucleus (16–18). Finally, NP is critically involved in the selective packaging of the viral genome segments – both directly through specific amino acid residues (19, 20) and indirectly though determining the accessibility of RNA structures important for packaging (21, 22).

We previously identified an NP substitution (NP:F346S) that was sufficient to significantly enhance the replication and transmissibility of the A/Puerto Rico/8/1834 (PR8) strain of IAV in guinea pigs while selectively decreasing the expression of the neuraminidase (NA) gene segment (23, 24). This finding suggested (a) that NP plays an unappreciated role in selectively regulating the expression of individual viral genes, and (b) that this mode of gene regulation may be involved in modulating transmission potential. Given that gene segment replication and transcription occur in the context of the vRNP, and that the vRNPs of all eight gene segments are thought to largely be structurally and functionally equivalent, it is not clear how substitutions in NP could result in selective modulation of NA segment expression.

Here, we dissect the mechanism by which specific residues in NP selectively modulate NA segment expression. In addition, we pinpoint the specific determinants within the NA genomic RNA that are required for susceptibility to selective regulation by NP. Altogether, these results illuminate a new mode of selective gene regulation by influenza viruses that may play an important role in host adaptation and transmission.

## Results

### NP:F346S suppresses NA segment replication but not mRNA transcription

We examined the effects of NP:F346S on NA vRNA abundance over the course of a single PR8 replication cycle. Similar to our previous findings, NP:F346S reduced NA expression nearly 20-fold by 12 hours post-infection (hpi) (24), while leaving HA expression largely unaffected (**Fig 1A**). We previously showed that NP:F346S also decreased NA mRNA abundance, raising the possibility that this substitution directly affected all NA segment-derived RNA species (24). To determine whether the effects of NP:F346S are specific for vRNA, we compared levels of primary NA mRNA synthesis between NP:WT and NP:F346S in the presence of 100µg/mL cycloheximide.

**Fig 1.**
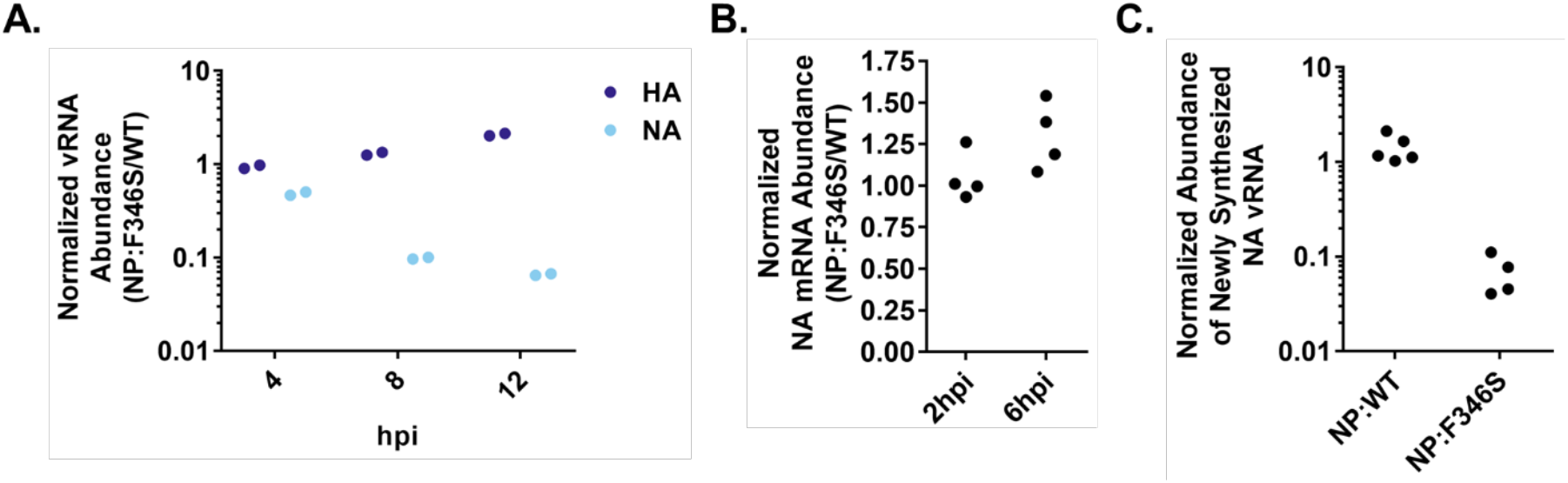
NP:F346S affects NA vRNA replication but not primary transcription. **A.)** Abundances of NA and HA vRNA (measured by RT-qPCR on total cellular RNA) at the indicated timepoints following infection of MDCK cells at an MOI of 0.1 NP-expressing units (NPEU)/cell under single cycle conditions. Data represent values obtained during infection with PR8-NP:F346S, normalized to values obtained during infection with PR8-NP:WT. The data shown are individual cell culture well replicates representative of the data obtained through two similar experiments. **B.)** NA mRNA abundances (measured by RT-qPCR on total cellular RNA) in PR8-NP:F346S-infected MDCK-SIAT1 cells, normalized to values obtained during infection with PR8-NP:WT. Infections were initiated at MOI=5 TCID_50_/cell in the presence of 100µg/mL cycloheximide. Data points indicate individual cell culture well replicates pooled from two independent experiments. **C.)** Abundances of newly synthesized NA vRNA in PR8-NP:F346S and PR8-NP:WT infected cells, as measured by 4-thiouridine (4SU) pulse labeling. MDCK cells were infected with PR8-NP:WT or PR8-NP:F346S at an MOI of 5 TCID_50_/cell for 7hrs, followed by 1hr pulse with 500µM 4SU. Cellular RNA was then harvested and the abundance of 4SU-labeled viral RNAs were determined by RT-qPCR using a universal, vRNA-sense specific primer for the RT reaction followed by segment-specific primers for the qPCR. Data points indicate individual cell culture well replicates pooled from two independent experiments.

Cycloheximide blocks translation of the viral replicase machinery needed for vRNA synthesis, thus only allowing primary transcription of viral mRNAs from incoming vRNPs (25). We observed no differences in the NA mRNA levels between WT and NP:F346S in the presence of cycloheximide, indicating that NP:F346S has no effects on primary mRNA transcription (**Fig 1B**).

To determine whether the effect of NP:F346S on NA vRNA abundance is due to reduced synthesis, as opposed to a decrease in stability, we pulsed infected cells with 4-thiouridine (4SU) for one hour, and measured the amount of vRNA synthesized during the pulse by performing RT-qPCR on 4SU-labeled RNAs using a universal, vRNA-specific primer for the RT reaction followed by segment-specific primers for the qPCR, or a tagged, vRNA and segment-specific primer for the RT reaction followed by a combination of a tag-specific and segment-specific primer for the qPCR step (**Figs 1C and S1 Fig.**). In both cases, levels of the 4SU-containing newly synthesized NA vRNA were over 10-fold lower during NP:F346S infection compared with NP:WT (**Figs 1C and S1 Fig.**). Altogether our data indicate that NP:F346S specifically affects the synthesis of new NA vRNA molecules during infection.

### The effect of NP:F346S on NA segment expression is strain-specific

Given that NP is thought to play the same role in the replication of all viral genome segments, how can substitutions in NP selectively reduce synthesis of the NA RNA while leaving the other segments largely unaffected? We hypothesized that this specificity must depend upon unique motifs present with the NA segment. To test this hypothesis, we introduced the NP:F346S substitution into a divergent IAV strain of the H3N2 subtype, A/Udorn/307/72 (Udorn), and examined whether it reduced Udorn NA segment expression similar to what was observed with PR8.

In the Udorn background, NP:F346S had no appreciable effect on NA segment expression, indicating that the effects of NP:F346S are virus-strain dependent (**Fig 2A**). To test whether this strain-specificity arises from the differences in NA segment (versus NP or the viral polymerase complex), we generated chimeric viruses encoding the Udorn HA and NA segments along with the remaining six segments from PR8. Introduction of the NP:F346S substitution into this chimeric virus similarly had no effect on NA expression, indicating that the effects of NP:F346S on NA expression depend upon the specific sequence of the NA segment (**Fig 2B**).

**Fig 2.**
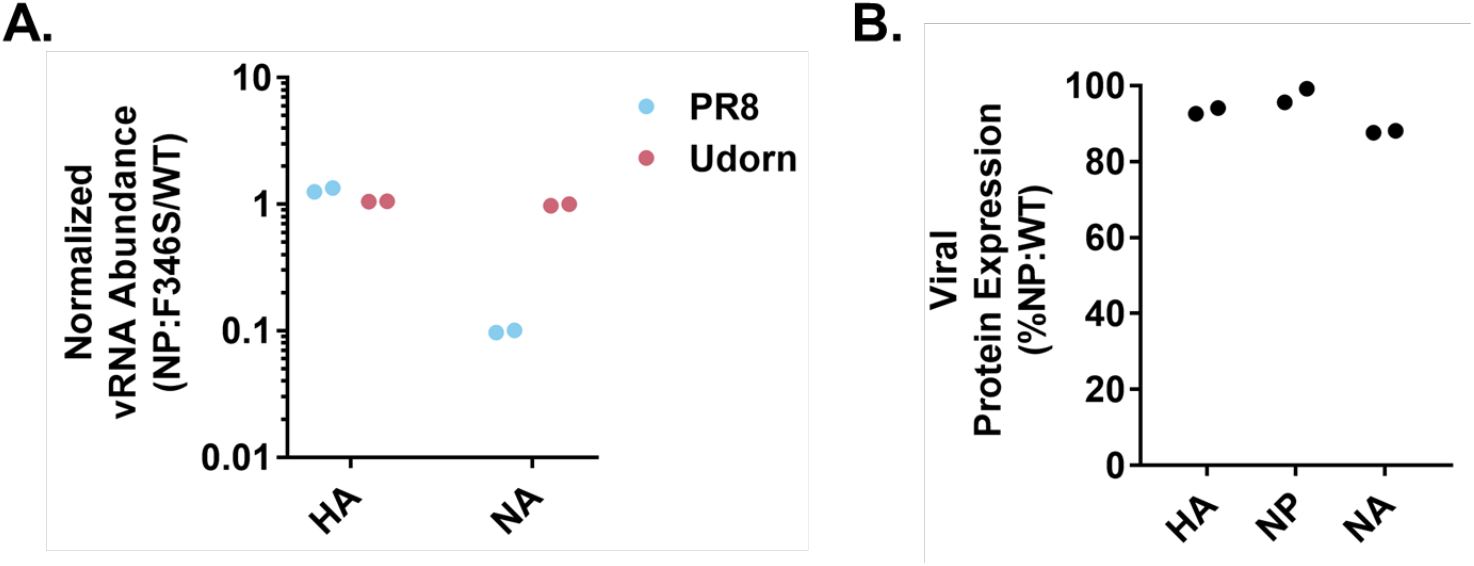
Susceptibility to the effects of NP:F346S is NA segment genotype specific. **A.)** Normalized vRNA abundances as determined by qRT-PCR in PR8-NP:F346S or Udorn-NP:F346S infected MDCK cells (MOI=0.1 NPEU/cell, 8hpi) expressed as fraction of PR8 NP:WT or Udorn NP:WT respectively. Secondary infection was blocked via the addition of ammonium chloride at 3hpi. The data points represent individual cell culture well replicates representative of the data obtained through two similar experiments. **B.)** Viral protein expression levels as determined by geometric mean fluorescence intensity (GMFI) in rPR8 Udorn HA/NA NP:F346S infected MDCK cells (MOI=0.03 TCID_50_/cell, 16hpi) expressed as a percentage of rPR8 Udorn HA/NA NP:WT. The data shown are individual cell culture well replicates representative of the data obtained through two similar experiments.

### Selective modulation of gene expression by NP:F346S depends upon segment UTR sequences

To pinpoint the specific motif(s) within the NA gene segment that confer susceptibility to selective modulation by NP:F346S, we first divided the NA segment into three broad functional regions: (a) the portion of the NA ORF that does not overlap known packaging signals, (b) the portions of the NA ORF that do overlap known packaging signals, and (c) the NA UTRs. We then tested each for their role in conferring sensitivity to the effects of NP:F346S.

To determine if any RNA sequence elements within the NA ORF (exclusive of the packaging signals as defined based on retention within defective interfering particles in a previous study (26)) were important for the effects of NP:F346S, we used the Codon Shuffle package (27) to introduce 227 silent substitutions within the region encompassing nucleotides 38-1319 of the NA segment while minimizing effects on codon frequency or di-nucleotide content (**Fig 3A**). The codon shuffled NA segment exhibited a similar decrease in its expression level as NA WT in the presence of NP:F346S, indicating the effects of NP:F346S on NA replication do not require motifs within the non-packaging signal region of the NA ORF (**Fig 3B**).

**Fig 3.**
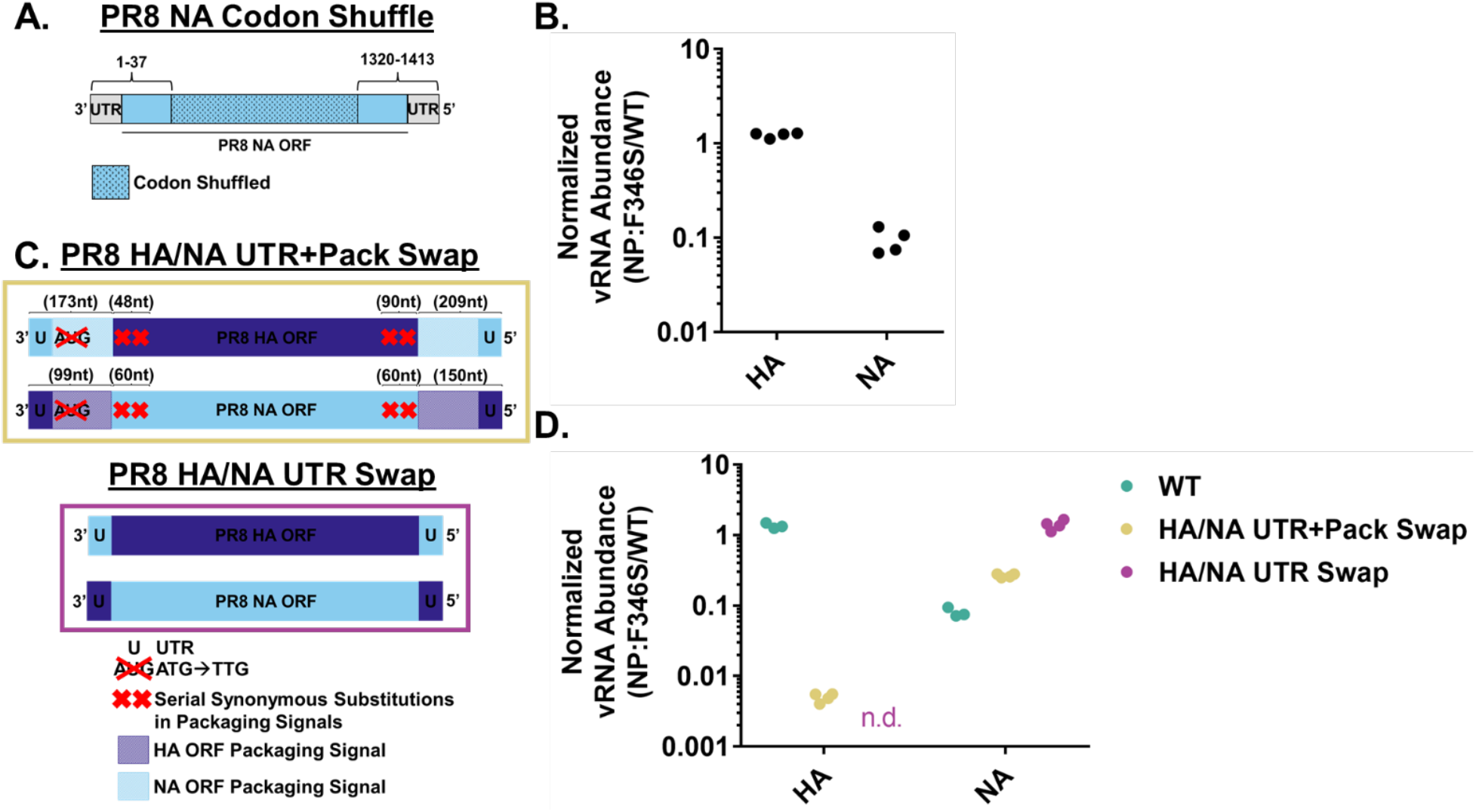
Susceptibility to NP-dependent regulation maps to the UTRs of the NA segment. **A.)** Schematic depiction of the codon shuffled PR8 NA construct. The Codon Shuffle program was used to introduce 227 silent mutations within the region encompassing nucleotides 38-1319 of the PR8 NA segment to alter features of the RNA sequence while minimizing changes in codon frequencies or dinucleotide content. **B.)** Relative abundances of HA and NA vRNA following infection of MDCK cells with the PR8 NA Codon Shuffle NP:F346S virus (MOI=0.1 NPEU/cell, 8hpi) as determined by RT-qPCR on cellular RNA expressed as a fraction of PR8 NA Codon Shuffle NP:WT respectively. Each data point represents a cell culture well replicate pooled from two separate experiments. **C.)** Schematic depictions of the PR8 HA/NA UTR+Pack Swap and PR8 HA/NA UTR Swap gene segments. The PR8 HA/NA UTR+Pack Swap segments were generated by replacing the UTRs and packaging signal regions of one segment (HA/NA) with those of the other segment (NA/HA). The start codon of the newly appended packaging signal for each segment was mutated to prevent the expression of any protein encoded by the packaging signal sequence. The packaging signals within the native ORFs were disrupted via the addition of silent substitutions to all codons to prevent duplication of the packaging signals in the swapped segments. The PR8 HA/NA UTR Swap gene segments were generated by swapping the UTRs of the PR8 HA/NA segments. **D.)** Relative abundances of the HA ORF containing or NA ORF containing segments from the PR8 NP:F346S, PR8 HA/NA UTR+Pack Swap NP:F346S, and PR8 HA/NA UTR Swap NP:F346S viruses in infected MDCK cells (MOI=0.1 NPEU/cell, 8hpi) as determined by RT-qPCR on total cellular RNA, expressed as a fraction of PR8 NP:WT, PR8 HA/NA UTR+Pack Swap NP:WT, and PR8 HA/NA UTR Swap NP:WT, respectively. N.d. indicates that the segment was below the limit of detection for the assay. Each data point represents a cell culture well replicate pooled from two separate experiments.

Not surprisingly, attempts to use codon shuffling to mutagenize the regions of the NA ORF that overlap the packaging signals resulted in non-viable viruses. As an alternative approach to examine the roles of the NA segment packaging signal regions and UTRs in determining sensitivity to NP:F346S, we generated two sets of recombinant PR8 viruses where we swapped terminal sequences between the HA segment (which is unaffected by NP:F346S) and the NA segment, and paired them with either NP:WT or NP:F346S (**Fig 3C**). One set of viruses contained chimeric HA-NA segments in which both the UTRs and packaging signals present within the terminal coding regions of the PR8 HA and NA segments were swapped (UTR+Pack swap) based on a previously described set of viable chimeric HA-NA segments (28). The other set of viruses contained segments in which only the UTRs of the PR8 HA and NA segments were swapped (UTR swap). For the UTR swap viruses, a segment encoding the HA ORF with the NA UTRs exhibited a severe packaging deficiency (**S2A Fig.**), and the abundance of the segment in infected cells was below the limit of detection for the qPCR assay.

We infected MDCK cells with these recombinant viruses and quantified the effects of NP:F346S on intracellular HA and NA vRNA levels (**Fig 3D**). Replacing the packaging signals and UTRs of the NA segment with those of the HA segment reduced the effect of NP:F346S on NA expression ∼3-fold (**Fig 3D**). Similarly, while WT HA expression is unaffected by NP:F346S, an HA segment containing the packaging signals and UTRs from the NA segment exhibited a >100-fold reduction in expression in the context of NP:F346S versus NP:WT (**Fig 3D**). Looking at the ratio of the HA and NA segments with the swapped UTRs and packaging signals in the viral stocks, the decrease in their abundance in the presence of NP:F346S corresponds to the observed expression decrease, suggesting that the observed changes in gene expression largely stem from changes in gene segment packaging ratios – likely a result of removing the packaging signals from their native context (**S2A/B Figs.**). Further, we observed that replacing the NA segment UTRs with those from HA completely eliminated the effect of NP:F346S on NA vRNA abundance (**Fig 3D**). Altogether, these data indicate that the selective effects of NP:F346S on vRNA synthesis depend upon the segment UTR sequences.

### Identification of specific nucleotides within the NA UTR that determine sensitivity to NP:F346S

We next sought to identify which specific elements within the NA UTR are required for susceptibility to modulation by NP:F346S. To do this, we took advantage of the high degree of similarity between the NA segment UTRs of PR8 (susceptible to the effects of NP:F346S) and Udorn (resistant to the effects of NP:F346S) (**Figs 2 and 4A**). The 3’ UTRs of the PR8 and Udorn NA segments differ in (a) the identity of the nucleotides at positions 4 and 13 (CC/UU for PR8/Udorn respectively) and (b) the sequence directly upstream of the initiating Met codon of the NA ORF (AAAUUU/ACUUC) for PR8/Udorn respectively) (**Fig 4A**). In the 5’ NA segment UTR, Udorn has a 9bp insertion relative to the PR8 sequence plus a few additional nucleotide substitutions (**Fig 4A**).

**Fig 4.**
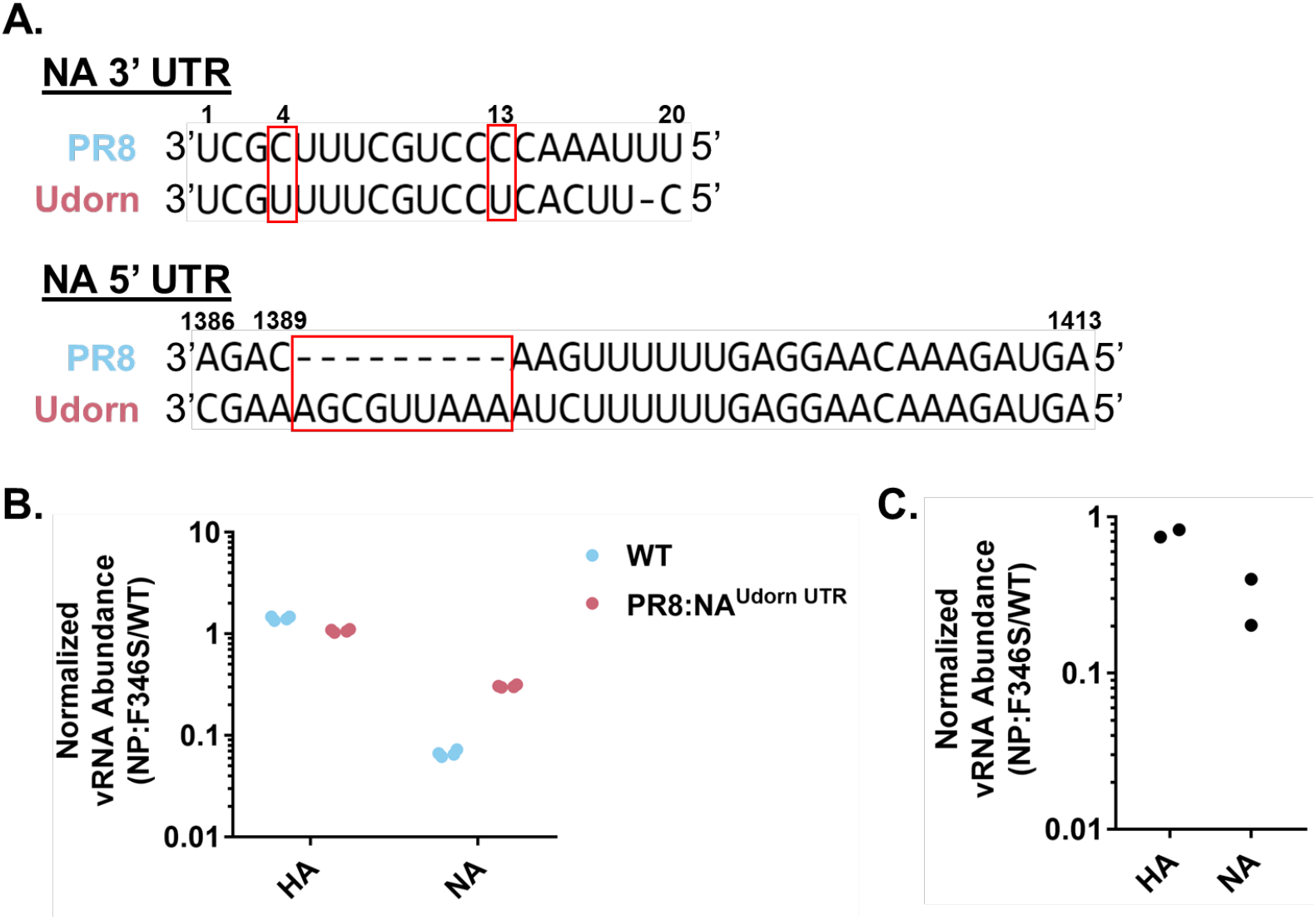
The UTRs of the Udorn NA segment confer resistance to regulation by NP:F346S. **A.)** Alignment of the PR8 NA and Udorn NA 3’ & 5’ UTRs using the M-Coffee alignment algorithm on the T-Coffee web server (29). Regions of interest are boxed in red. PR8 NA nucleotide numbering is shown. **B.)** Relative abundances of the HA and NA segments in MDCK cells infected with the PR8 NP:F346S, PR8:NA^Udorn UTR^ NP:F346S (MOI=0.1 NPEU/cell, 8hpi) viruses as determined by RT-qPCR expressed as a fraction of PR8 NP:WT and PR8:NA^Udorn UTR^ NP:WT respectively. Each data point represents a cell culture well replicate pooled from two independent experiments. **C.)** Relative abundances of the HA and NA segments in MDCK cells infected with PR8:Udorn HA, NA^PR8 UTR^ NP:F346S (MOI=0.03 NPEU/cell, 8hpi) virus as determined by RT-qPCR expressed as a fraction of PR8:Udorn HA, NA^PR8 UTR^ NP:WT. Each data point represents a cell culture well replicate from a single experiment.

We first confirmed that the difference in susceptibility of the PR8 and Udorn NA segments to the effects of NP:F346S is associated with the UTR sequences. We generated a virus in which the UTRs of the PR8 NA segment were replaced with those from the NA segment of Udorn (PR8:NA^Udorn UTR^) (**Figs 4A and 5**). The effects of NP:F346S on PR8:NA^Udorn UTR^ were reduced compared with WT PR8 NA, again indicating that the segment UTR sequences play a significant role in determining the segment specificity of the effects of NP:F346S on gene expression (**Fig 4B**). We also attempted to generate viruses where the UTRs of the Udorn NA segment were replaced with those from PR8, however, we were unable to rescue a virus with this chimeric Udorn/PR8 NA segment and NP:F346S. By replacing the internal gene segments of Udorn with those of PR8, we were able to rescue viruses containing a segment with the Udorn NA ORF and PR8 UTRs and NP:WT/F346S (PR8:Udorn HA,NA^PR8 UTR^). The viruses were still highly attenuated, reaching titers of only 10^4^-10^5^ infectious particles/mL. Replacing the Udorn NA UTRs with those of PR8 made it susceptible to the effects of NP:F346S, although to a lesser degree than PR8 NA (∼30% v. 5-10% of NP:WT respectively), further substantiating the role of the NA UTRs in regulation by NP:F346S (**Fig 4C**). Additionally, while the Udorn NA segment paired with a PR8 backbone exhibited no apparent defects in genome packaging, the Udorn NA:^PR8 UTR^ segment in a PR8 backbone did exhibit decreased packaging efficiency in the presence of NP:F346S, suggesting that some of the observed decrease in expression levels within infected cells might be due to decreases in delivered NA gene dose due to decreased packaging efficiency of the Udorn NA:^PR8 UTR^ segment (**S2C Fig.**).

We next generated a panel of recombinant viruses with chimeric PR8-Udorn NA UTR sequences (**Fig 5**).

**Fig 5.**
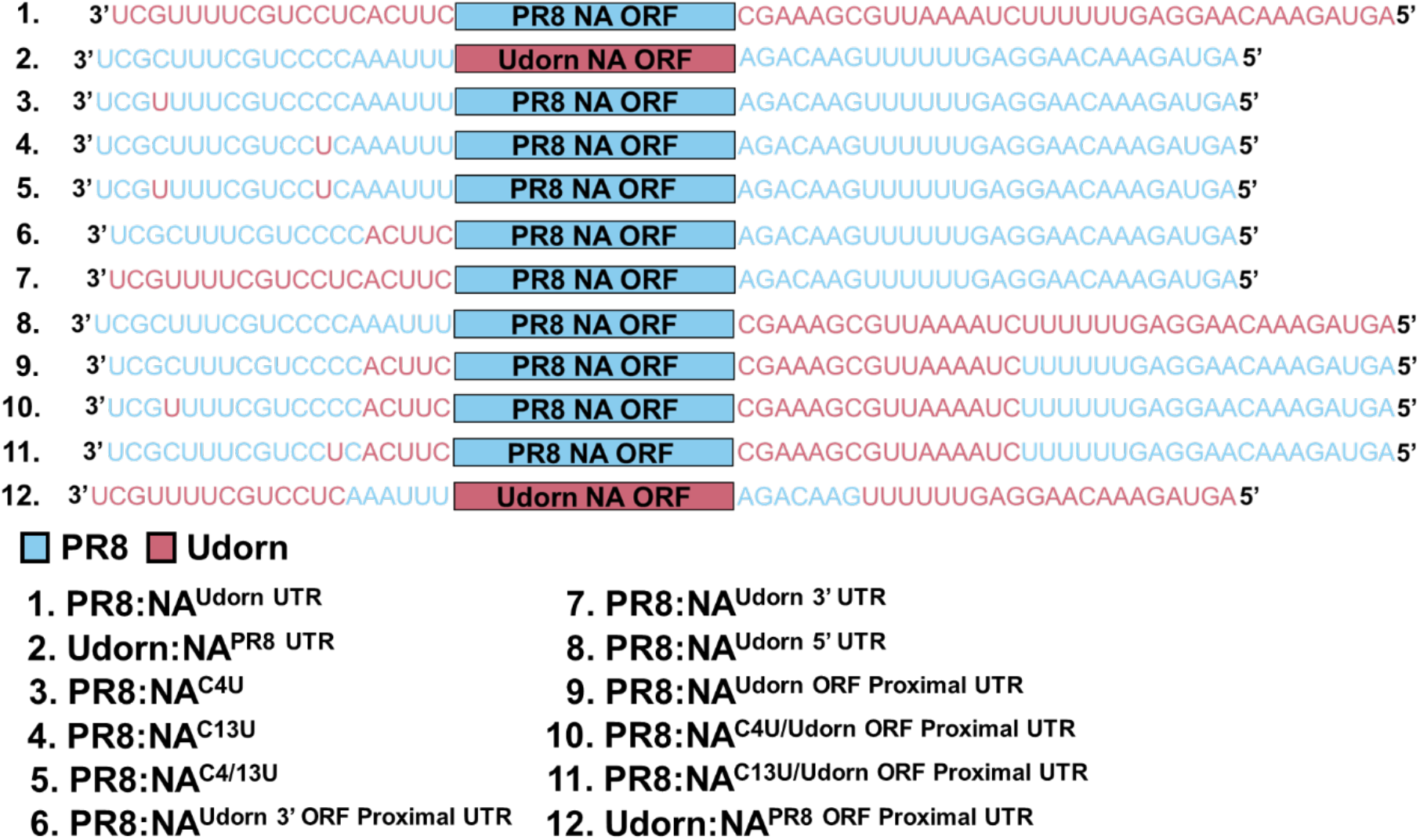
UTR sequences of the PR8-Udorn NA UTR chimeric constructs. The sequences derived from PR8 and Udorn NA are colored blue and pink, respectively. Sequences shown in negative sense, 3’->5’.

We first examined both the polymorphisms at positions 4 and 13 of the promoter and promoter proximal region in the 3’ UTR. Introducing individual C4U or C13U substitutions into the PR8 NA UTR slightly reduced the extent to which NP:F346S decreased NA vRNA levels (**Fig 6A**). Introducing both C4U and C13U together decreased the relative effect of NP:F346S on NA vRNA levels by roughly 6x (**Fig 6A**). Retaining the C’s at positions 4 and 13 while swapping the ORF proximal region of the PR8 NA 3’ UTR with that of Udorn (PR8:NA^Udorn 3’ ORF Proximal UTR^) slightly reduced the extent to which NP:F346S decreased NA vRNA levels, to an extent comparable with the individual C4U or C13U mutations (**Fig 6A**). Replacing the entire PR8 NA 3’ UTR with that of Udorn (PR8:NA^Udorn 3’ UTR^) (Has U4/13 and Udorn ORF Proximal UTR sequence) decreased the relative effect of NP:F346S to approximately the same level as the dual C4/13U mutations (**Fig 6A**). Taken together, these data demonstrate that sensitivity to the effects of NP:F346S is largely determined by the identity of nucleotides 4 and 13 in the 3’ UTR, however, additional motifs within the segment also likely contribute.

**Fig 6.**
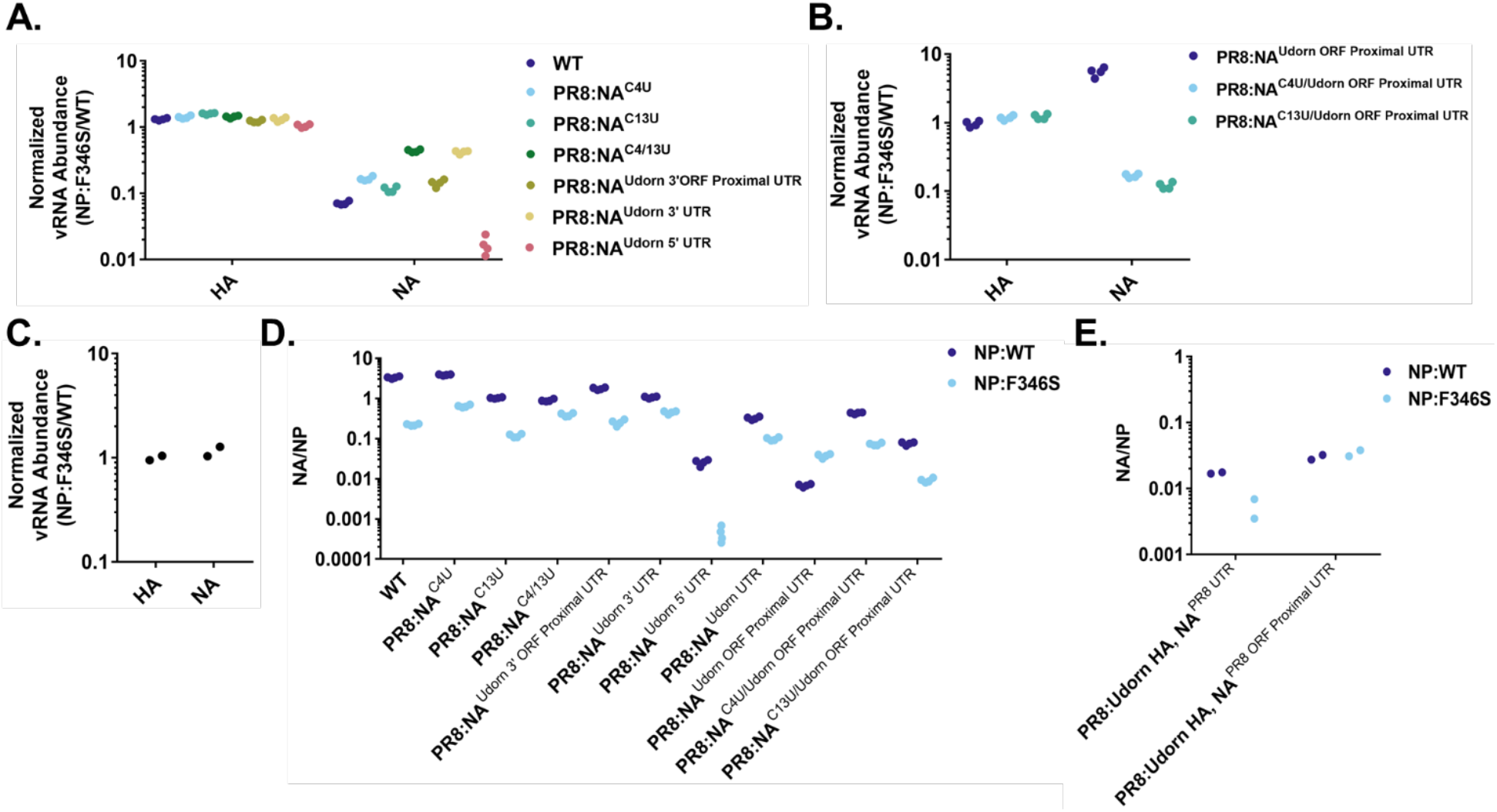
The effect of mutations in the PR8 NA UTRs on baseline expression levels and sensitivity to NP:F346S. **A.)** Relative abundances of the HA and NA segments at 8hpi in MDCK cells infected with the indicated viruses encoding NP:F346S at MOI=0.1 NPEU/cell, as determined by qRT-PCR normalized to the NP:WT-encoding versions of the same viruses. Each data point represents an individual cell culture well replicate pooled from two independent experiments. **B.)** Relative abundances of the HA and NA segments in MDCK cells infected with the indicated viruses encoding NP:F346S (MOI=0.1 NPEU/cell, 8hpi), as determined by qRT-PCR normalized to the NP:WT-encoding versions of the same viruses. Each data point represents an individual cell culture well replicate pooled from two independent experiments. **C.)** Relative abundances of the HA and NA segments in MDCK cells infected with the PR8:Udorn HA, NA^PR8 ORF Proximal UTR^ NP:F346S virus (MOI=0.03 NPEU/cell, 8hpi) as determined by qRT-PCR normalized to PR8:Udorn HA, NA^PR8 ORF Proximal UTR^ NP:WT. Each data point represents an individual cell culture well replicate from a single experiment. **D,E.)** Data from experiments shown in **(4B and 6A,B)** and **(4C/6C)** respectively, showing the intracellular abundances of the indicated chimeric NA segment vRNAs normalized to NP vRNA levels (in the context of NP:WT or NP:F346S) in infected MDCK cells (MOI= 0.1 **(D)** or 0.03 **(E)** NPEU/cell 8hpi) as determined by qRT-PCR on total cellular RNA. The data represents two cell culture well replicates pooled from either two independent experiments **(D)** or a single experiment **(E)**.

We next examined the roles of the ORF proximal regions of the NA UTRs. As described above, just replacing the 3’ PR8 NA ORF proximal region with that of Udorn (PR8:NA^Udorn 3’ ORF Proximal UTR^) did not have much of an effect of susceptibility to NP:F346S (**Fig 6A**). Replacing the 5’ UTR of the PR8 NA with that of Udorn (PR8:NA^Udorn 5’ UTR^) enhanced susceptibility to NP:F346S by ∼4x (**Fig 6A**) likely due to the fact that there was an additional packaging defect for the segment in the presence of NP:F346S (**S2D Fig.**). Interestingly, replacing both the ORF proximal regions of the PR8 NA with those from Udorn (PR8:NA^Udorn ORF Proximal UTR^) (Has 4/13C and Udorn NA ORF proximal sequences) made the segment resistant to NP:F346S, actually increasing expression ∼5-6x relative to NP:WT (**Fig 6B**).

The only differences between the PR8:NA^Udorn UTR^ segment, which was partially resistant to the effects of NP:F346S (**Fig 4**), and PR8:NA^Udorn ORF Proximal UTR^, which was completely resistant, were the identity of nucleotides 4 and 13 of the 3’ UTR (**Fig 4**), so we next asked whether these nucleotides were responsible for the resistance phenotype observed for PR8:NA^Udorn ORF Proximal UTR^). Mutating the C at position 4 or 13 of the PR8:NA^Udorn ORF Proximal UTR^) segment to U (PR8:NA^C4U/Udorn ORF Proximal UTR^/PR8:NA^C13U/Udorn ORF Proximal UTR^), restored the susceptibility of the segment to NP-dependent regulation, again emphasizing the importance of positions 4 and 13 of the PR8 NA 3’ UTR to determining the effects of NP:F346S (**Fig 6B**). Interestingly, we also found that replacing the ORF proximal regions of the Udorn NA UTR with those of PR8 (PR8: Udorn HA,NA^PR8 ORF Proximal UTR^) (Has 4/13U and PR8 NA ORF Proximal Sequences) made the segment resistant to NP:F346S (**Fig 6C**). In conclusion, susceptibility of a gene segment to the effects of NP:F346S depends upon a specific combination of nucleotide identities at positions 4 and 13 of the 3’ UTR and ORF proximal sequences (4/13C and Udorn NA ORF proximal UTRs, or 4/13U and PR8 NA ORF proximal UTRs).

As several of the PR8-Udorn NA UTR mutant viruses were highly attenuated, we wanted to determine whether there were any compensatory mutations that may have emerged that could potentially confound our results. We performed next-generation sequencing on these viruses and found that the only virus with any mutations over ∼30% in the population was PR8:NA^Udorn UTR^ NP:F346S, which had a fixed nonsynonymous substitution in PB2 (E191G). We cannot rule out the possibility that this mutation affects NA segment expression.

We hypothesized that the effects of different UTR mutations on sensitivity of NA expression levels to NP:F346S could arise from two distinct mechanisms: (1) abrogating the selective effect of NP:F346S on NA vRNA synthesis, restoring NA vRNA levels to those observed in the context of NP:WT, or (2) reducing NA levels in the context of NP:WT, bringing them closer to what is observed with NP:F346S. To distinguish between these possibilities, we compared the expression of the different NA UTR mutant constructs to NP vRNA levels (which are unaffected by the NP:F346S substitution) (**Figs 6D and 6E**). We observed that the expression of all the PR8/Udorn NA UTR chimeric constructs except for PR8:NA^C4U^ was reduced compared to WT NA in the context of NP:WT, and in some cases, lower than the level observed for the WT NA segment in the presence of NP:F346S (**Figs 6D and 6E**). Thus, none of the UTR mutants tested mitigated the effects of NP:F346S by simply restoring NA levels to those observed in the context of NP:WT. The second possibility, that these mutations appeared to reduce sensitivity to NP:F346S because they reduced NA levels in the context of NP:WT to levels associated with NP:F346S, was also not supported by these data. For instance, the PR8:NA^Udorn ORF Proximal UTR^ segment exhibited increased expression in the presence of NP:F346S, while the PR8:NA^Udorn 5’ UTR^ segment exhibited an even more substantial decrease in the presence of NP:F346S than the WT NA (**Fig 6D**). Altogether, our data suggest that it is impossible to cleanly separate the effects of the UTR sequences on susceptibility to the effects of NP:F346S from their broader effects on baseline expression levels in the context of NP:WT.

### NP:F346S has no measurable effects on NP RNA binding or oligomerization

While the UTR sequences of the NA segment are clearly involved in determining the segment-specificity of the effects of NP:F346S on gene expression, the specific mechanisms involved are not obvious. NP:F346S is not located in any previously described functional domains, thus is was not immediately apparent how the NP:F346S substitution might alter NP protein function (12,14,30–34).

NP has two well-described biochemical activities that are required for the synthesis of full-length viral RNA transcripts: RNA binding and oligomerization (12,15,35). To determine whether F346S affects the RNA-binding activity of NP, we compared the *in vitro* RNA binding affinities of purified C-terminal his-tagged versions of the NP:WT and NP:F346S proteins using bio-layer interferometry (BLI) (**Fig 7A**). NP:F346S was associated with a slightly higher K_D_ compared with NP:WT (3.216±0.1088nM vs. 2.477±0.09097nM), however, it was not clear that this difference was biologically significant (**Fig 7A**). Although these data suggest that the F346S substitution has minimal effects on the RNA-binding affinity of NP, our *in vitro* assay may have failed to fully recapitulate conditions as they occur during infection.

**Fig 7.**
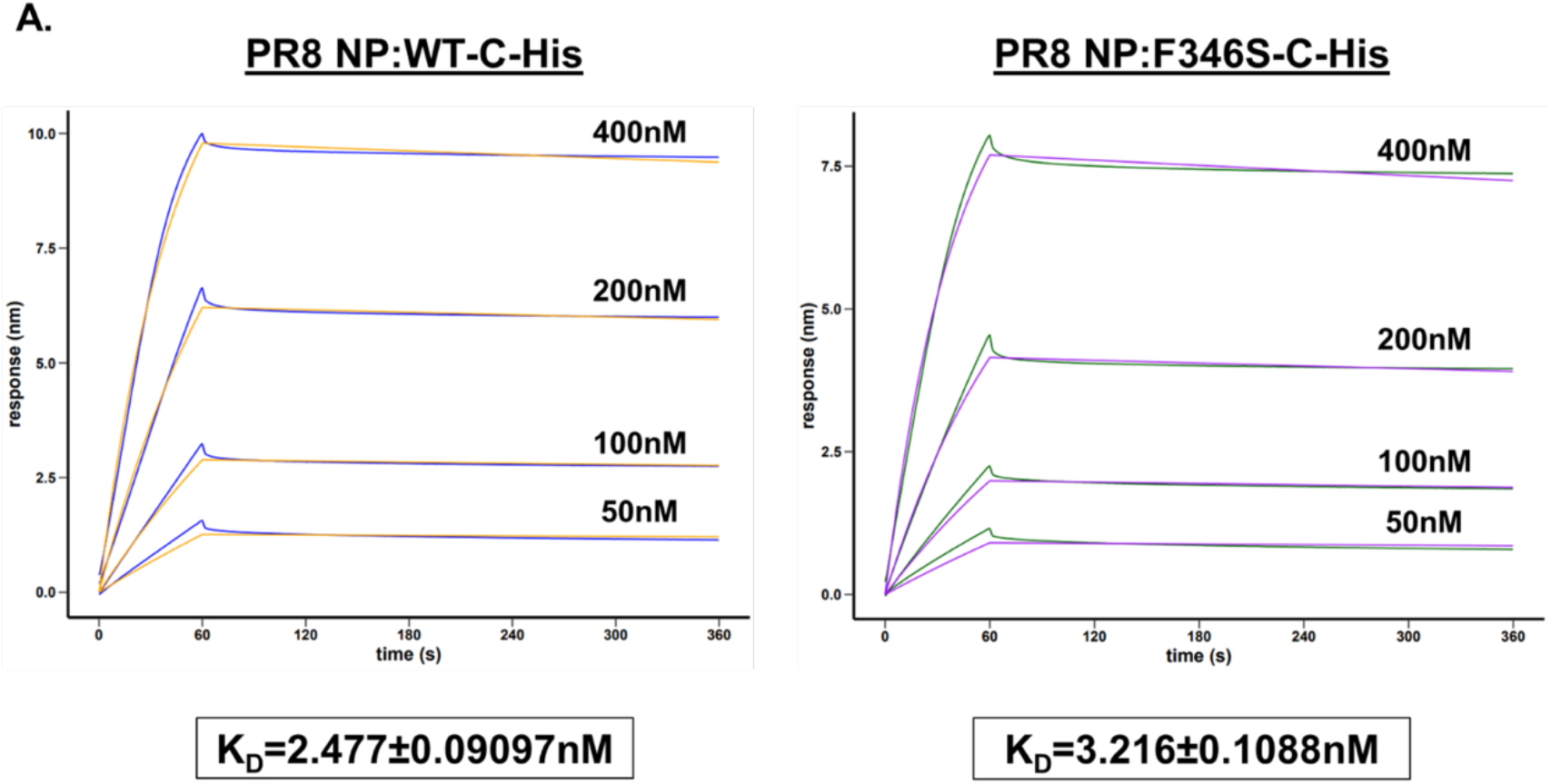
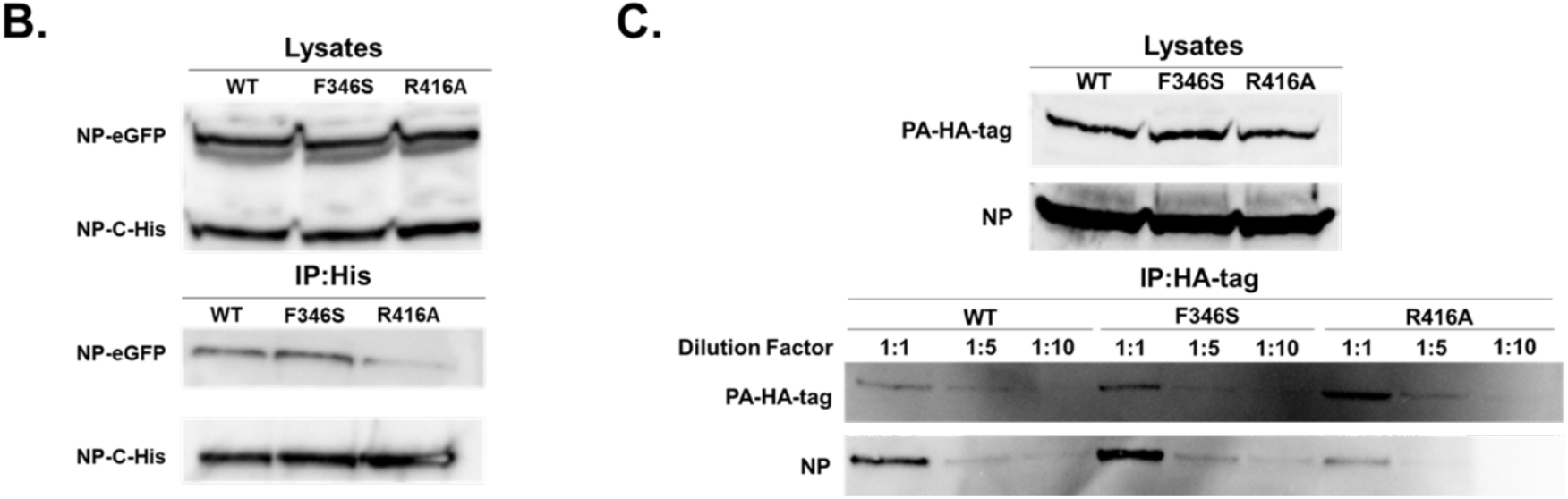
NP:F346S does not affect NP RNA binding or oligomerization. **A.)** RNA binding kinetics of the PR8 NP:WT-C-His and PR8 NP:F346S-C-His proteins as determined by BLI. The raw data is colored blue/green and the fitted data is colored orange/purple for the NP:WT/F346S-C-His proteins, respectively. **B.)** Co-immunoprecipitation (IP) of eGFP and His-tagged versions of the indicated NP proteins. 293T cells were transfected with expression vectors encoding the eGFP-and His-tagged versions of either WT, F346S, or R416A NP proteins. Lysates were harvested after 24hrs. His-tagged NP was immunoprecipitated, and then IP samples were probed via western blot with anti-eGFP and anti-6x His antibodies. Western blots of total cell lysates stained with an anti-NP antibody are also shown. **C.)** Co-IP of vRNP-associated NP and PA. Cells were transfected with plasmids encoding the vRNP complex (PB2, PB1, PA-HA-tag, and NP (WT, F346S, or R416A) and a vRNA template (NA vRNA). Lysates were harvested 24hrs post transfection, and vRNP complexes were IP-ed using an anti-HA-tag antibody. Undiluted, 1:5 diluted, or 1:10 diluted IP-ed protein was probed with anti-NP and anti-HA-tag antibodies via western blot. Western blots of whole cell lysates shown for comparison.

We next evaluated whether NP:F346S affects the oligomerization of NP monomers. We overexpressed both His-tagged and eGFP-tagged versions of either NP:WT or NP:F346S in 293T cells, and quantified the amount of eGFP-NP that co-immunoprecipitated with His-NP by western blot. As a positive control, we assessed the effect of the oligomerization-deficient R416A mutant (12,14,36,37) in our assay and measured a substantial reduction in pull-down efficiency (**Fig 7B**). In contrast, we did not observe any effect of F346S on the co-immunoprecipitation efficiencies of His-NP and eGFP-NP, suggesting that NP:F346S does not significantly affect the ability of NP to oligomerize, at least under *in vitro* over-expression conditions (**Fig 7B**).

Additionally, we evaluated whether NP:F346S decreases the NP content of vRNPs. We overexpressed the PB2, PB1, HA-tagged PA, NP (WT/F346S/R416A) proteins and a NA vRNA template in 293T cells to generate vRNPs and visualized the amount of NP that co-immunoprecipitated with HA-tagged PA via western blot. Mirroring our data looking at NP monomer association, there was a substantial reduction in pull-down efficiency for the oligomerization-deficient NP:R416A mutant, but no difference in the pull-down efficiencies between NP:WT and F346S (**Fig 7C**). These data suggest that the NP content of vRNPs is not affected by NP:F346S.

Altogether, these data suggest that F346S has minimal effects on the RNA-binding and oligomerization activities of NP, at least in *in vitro* binding assays. If true in the context of infection, it would suggest that the effects of this substitution on NA segment replication occur through some other, uncharacterized feature of NP protein biology.

### A cluster of aromatic residues within NP governs NA segment replication

Finally, we examined the effects of alternative substitutions at the NP:F346 locus on NA expression. Introducing a silent substitution (T1082C) into the NP:F346 codon had no effects on NA protein expression levels, indicating that the effects of NP:F346S require the amino acid substitution (**Fig 8A**). Interestingly, we observed that any amino acid substitution at position 346 resulted in a selective decrease in NA expression indicating that a phenylalanine is required at NP position 346 for maximal NA expression (**Fig 8A**). To better understand the need for a phenylalanine residue at this position, we examined the surrounding protein structure. We noticed two additional aromatic residues (Y385 & F479) directly adjacent to F346 that could potentially interact via π-π stacking interactions (**Fig 8B**). Mutation of either Y385 or F479 to alanine resulted in a selective decrease in NA abundance, though not quite as pronounced as that observed for NP:F346S (**Fig 8C**). These data suggest that maximal expression of the NA gene segment depends upon a cluster of aromatic residues F346, Y385, and F479 in NP.

**Fig 8.**
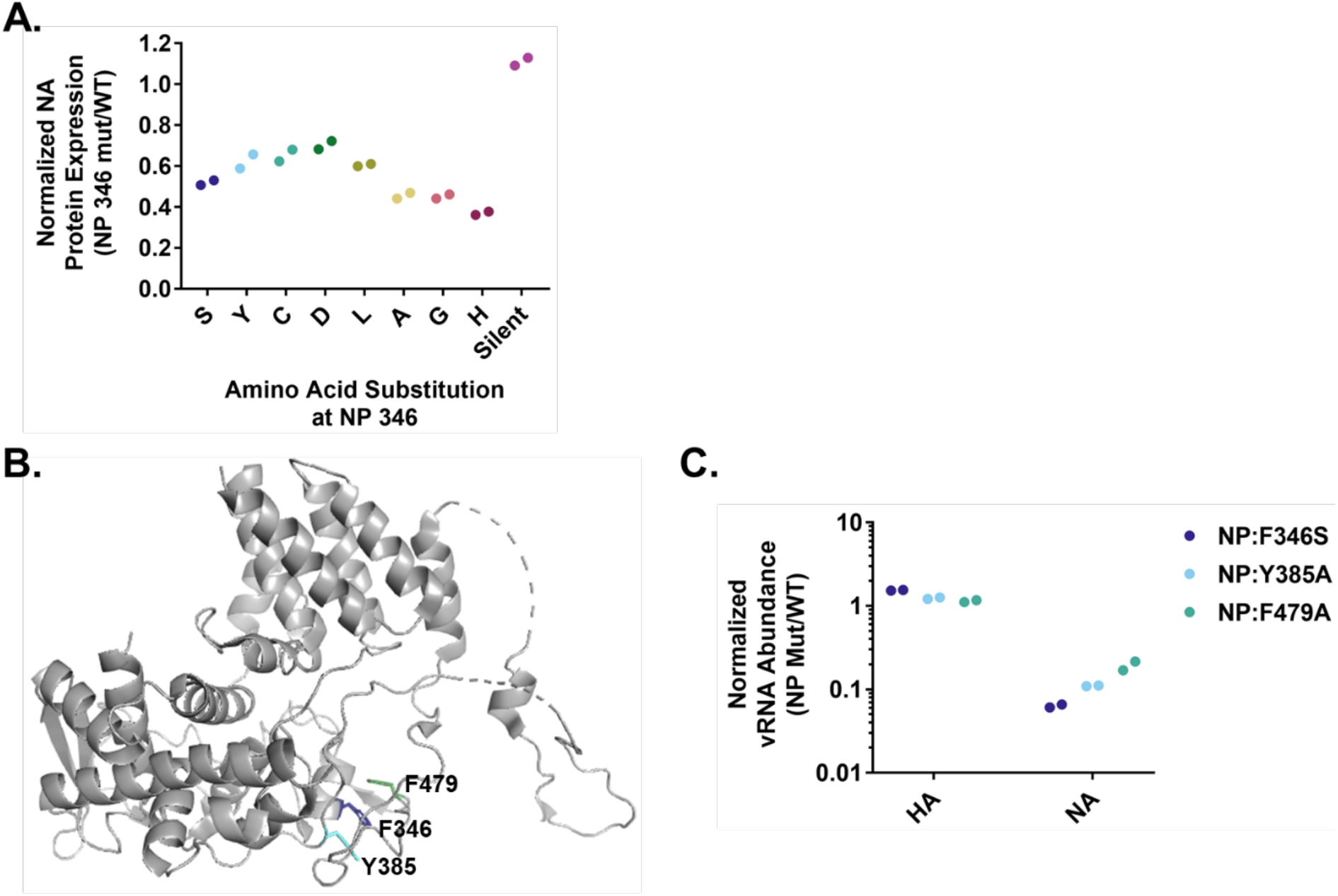
A cluster of aromatic residues is involved in the regulation of NA gene segment expression. **A.)** Normalized NA protein expression levels in cells infected with the indicated PR8 NP 346 mutant viruses (MOI=0.03 TCID_50_/cell, 16hpi) as determined by geometric mean fluorescent intensity (GMFI) expressed as a fraction of PR8 NP:WT. The data shown are individual cell culture well replicates representative of the data obtained through two similar experiments. **B.)** Location of F346, Y385, and F479 in the NP protein visualized using the PyMol software (PDB 2IQH). **C.)** Normalized viral RNA abundance in PR8 NP:F346S, PR8 NP:Y385A and PR8 NP:F479A infected MDCK cells (MOI=0.1 TCID_50_/cell, 8hpi) as determined by RT-qPCR and expressed as a fraction of PR8 NP:WT. The data shown are individual cell culture well replicates representative of the data obtained through two similar experiments.

## Discussion

Our results describe a surprising role for NP in the selective regulation of NA segment synthesis during IAV infection. We found that substitutions at NP:F346 can specifically decrease the rate of NA vRNA synthesis while leaving the other gene segments largely unaffected. The specificity of this effect largely depends upon specific sequence motifs within the NA segment UTRs, demonstrating how interactions between NP and the individual gene segment UTRs can selectively modulate gene segment replication and expression.

Our results raise several additional questions, one of which concerns the role of the F346-Y385-F479 motif in NP function. The F346/Y385/F479 residues are highly conserved among IAV NP genes (38), indicating the importance of this motif for viral fitness in humans. NP promotes vRNA replication by stabilizing the positive sense cRNA replicative intermediate and by acting as an elongation factor (15, 25). Previous studies have demonstrated that these functions require both the RNA binding and oligomerization activities of NP (15,25,35). Surprisingly, we found that NP:F346S does not appreciably affect the RNA binding or oligomerization activities of NP *in vitro* (**Fig 7**), however, more in depth studies examining NP assembly and recruitment to vRNPs in the context of infection would aid in further substantiating whether NP:F346S affects these activities. If NP-RNA binding and oligomerization are not affected by NP:F346S, this raises the question of how substitutions at NP:F346 can modulate vRNA replication kinetics. One possibility is that positions NP F346-Y385-F479 govern interactions with other viral and/or cellular proteins involved in IAV gene segment replication (33,34,39– 46). Another intriguing possibility is that substitutions at NP positions F346-Y385-F479 affect the types of specific viral RNA species that are produced, such as svRNAs, which can modulate replication in a segment-specific manner (47, 48). Finally, based on the NP structure, π-π stacking interactions between these residues may stabilize the structure of the loop regions where Y385 and F479 are located (**Fig 8**), thus these residues may play a role in maintaining the structural integrity and stability of NP.

The effects of the NP:F346S substitution clearly depend upon specific sequence motifs in the NA segment UTRs, however, this relationship is complicated. We identified two regions in the NA UTR that were important for determining both susceptibility to NP-dependent regulation and baseline NA segment expression levels in the context of WT NP: nucleotides 4 and 13 in the promoter/extended duplex region and the 3’ and 5’ ORF proximal regions. The NA segment that demonstrated the lowest sensitivity to the effects of NP:F346S was one that contained a combination of UTR features from PR8 and Udorn: C’s at positions 4 and 13 (as in WT PR8) and the Udorn-derived 3’ & 5’ ORF proximal sequences (**Figs 5 and 6B**). Interestingly, this specific NA segment also exhibited a >10-fold reduction in baseline NA expression in the absence of NP:F346S (**Fig 6D**). Disrupting this pairing by mutating one of the C’s at position 4 or 13, or replacing one of the Udorn-derived ORF proximal regions with that from PR8 restored the effects of NP:F346S on NA synthesis (**Figs 5 and 6A,B**). All NA constructs that we tested that were less sensitive to the effects of NP:F346S, with the exception of PR8:NA^C4U^, also exhibited significantly lower baseline levels of NA expression under WT NP conditions, indicating that sensitivity to the effects of NP:F346S cannot be uncoupled from baseline NA expression levels.

For all eight genome segments, gene expression is influenced by the structure of the viral promoter that is formed by base-paring interactions between the 3’ and 5’ UTRs (49). Base-pairing between positions 4 of the 3’ and 5’ UTR and between positions 13 of the 3’ UTR and 14 of the 5’ UTR influence the promoter structure and stability and have been shown to be important for regulating gene segment replication and transcription (6,49–52). For the NA segment, a C at position 4 in the 3’ UTR (which is unable to base-pair with the A at position 4 of the 5’ UTR) promotes genome replication, while a U at this position favors mRNA transcription (50). For PR8, NA is the only segment with C’s at both positions 4 and 13 of the 3’ UTR, and thus has the fewest number of base-pairing interactions based on the traditional panhandle structure of the IAV promoter. This unique feature may make the NA segment of PR8 uniquely dependent on WT NP to facilitate the stable interaction between the promoter and the viral replicase.

The ORF-proximal regions of the UTRs also influence gene segment expression, however, the exact mechanism(s) remain unclear (5,7–10). Consistent with this, we observed that the ORF proximal regions of the NA UTRs play important roles in both regulating the baseline expression level of the NA segment and in determining sensitivity to NP:F346S. Relative to Udorn and the remaining seven gene segments of PR8, the PR8 NA segment UTRs harbor a unique extended stretch of base-pairing from 3’-nt14/5’-nt15 to 3’-nt17/5’-nt18 located within the poly U stretch of the 5’ UTR. Given the hypothesized role for NP in promoter escape and elongation (15), the NA segment may be particularly dependent on WT NP (and thus sensitive to NP:F346S) to allow the polymerase to bypass this extended base-pairing region during elongation of the nascent vRNA. Altogether, our data suggest that the unique sequence and presumed structure of the PR8 NA segment UTRs confer elevated sensitivity to perturbations of NP function and thus, likely explain the segment specificity of the effects of NP:F346S.

Through mutations in the UTR sequences of individual segments, IAV can more finely coordinate the expression of the eight individual gene segments without altering the protein coding capacity of the segments or polymerase activity. Similar to our findings, a recent study demonstrated that the 3’ UTR of the HA segment played a role in regulating HA expression in a segment-specific manner, and that this regulation was only observed when the HA segment had to compete with the remaining seven segments for replication/transcription (7). An additional study using a reporter system also found that the UTR sequences of the segments affected the ability of the segments to compete with one another for access to the viral polymerase (52). Altogether, these studies highlight the importance of the individual segment UTR sequences in maintaining the optimal balance in expression of the eight gene segments during infection. Our results further demonstrate how slight perturbations in polymerase or NP function can affect the expression of specific segments to a greater degree than others.

What are the implications of NA (and potentially other viral gene segments) being sensitive to individual substitutions in NP? HA and NA facilitate viral particle attachment and release respectively, and balancing these activities is necessary for maintaining viral fitness (53–62). HA and NA evolve at faster rates than the rest of the IAV genome due to immune selection (63). Immune escape substitutions within HA and/or NA often alter glycoprotein function and require compensatory mutations to restore their functional balance and viral fitness (64–72). If substitutions in NP tune NA expression, it expands the number of available genetic pathways maintaining HA/NA functional balance. The functional link between the NP and NA segments that we establish here also has important consequences for reassortment, as the need to maintain compatible NP and NA genotypes may constrain the repertoire of viable reassortant progeny when heterologous viral strains mix. Finally, variation in NP-requirements between segments could influence patterns of expression kinetics, as the concentration of NP within the cell is dynamic over time. Altogether, our results highlight the potential of genome segmentation to facilitate dynamic changes in gene expression patterns through mechanisms that are not readily available to non-segmented viruses. This regulatory agility may help promote viral adaptation in response to changing host environments.

In summary, we identified a novel mechanism through which interactions between NP and other gene segment UTRs facilitate selectively regulation of viral gene expression. Our data reveal a new functional domain in the NP protein and suggest a broader role for NP in selective regulation of individual viral gene segments. These findings demonstrate how the expression of individual gene segments can be modulated to maximize viral fitness under different host conditions.

## Materials and Methods

### Plasmids

The A/Puerto Rico/8/34 and A/Udorn/72 reverse genetics plasmids were gifts from Drs. Adolfo Garcia-Sastre and Kanta Subbarao, respectively. The pCI vector was provided by Dr. Joanna Shisler. The lentivirus generation plasmids-pHAGE2-EF1aInt WSN HA W, HDM Hgpm2, HDM tatlb, pRC CMV Rev1b, HDM VSV-G were provided by Dr. Jesse Bloom. The peGFP-C1 plasmid for generating C-terminal eGFP-tagged proteins was provided by Dr. Andrew Mehle.

Point mutations were introduced into the PR8 NP segment via site-directed mutagenesis and *Lgu*I restriction sites were added to both ends. The inserts were then digested with *Lgu*I, ligated into the pDZ vector, and transformed into chemically competent *E. coli* cells via the heat-shock method. Insert sequences were confirmed via sanger sequencing.

PR8 NA ORF HA UTR+Pack Swap and PR8 HA ORF NA UTR+Pack inserts were generated via overlap extension PCR with primers designed to introduce the PR8 HA or PR8 NA UTR+Packaging Signal regions at the ends of the PR8 NA ORF or PR8 HA ORF using the pDZ PR8 NA or pDZ PR8 HA plasmid as a template respectively. Primers were used to add *Lgu*I restriction sites to each end of the inserts. PR8 NA ORF HA UTR and PR8 HA ORF NA UTR inserts were generated via PCR with primers designed to add the PR8 HA or PR8 NA UTRs to the PR8 NA or HA ORFs using the pDZ PR8 NA or pDZ PR8 HA plasmid as a template respectively. *Lgu*I restriction sites were added to each end. The inserts were then digested with *Lgu*I, ligated into the pDZ vector, and transformed into chemically competent *E. coli* cells via the heat-shock method. Insert sequences were confirmed via sanger sequencing.

For UTR chimera constructs, inserts were generated via PCR with primers designed to introduce the 4U and/or 13U mutations into the PR8 NA UTR, or to replace specific regions of the PR8 NA UTR at the 3’ and/or 5’ ends or the Udorn NA UTR at the 3’ and/or 5’ ends with the equivalent region(s) present in the Udorn NA UTR or PR8 NA UTR respectively. For the plasmids containing a chimeric Udorn-PR8 NA segment with the Udorn NA ORF, the A763C silent mutation was introduced to the Udorn NA sequence to disrupt an internal *Lgu*I restriction site. *Lgu*I restriction sites were added to the ends of the chimeric segments via PCR, the inserts were then digested with *Lgu*I, ligated into the pDZ vector, and transformed into chemically competent *E. coli* cells via the heat-shock method. Insert sequences were confirmed via sanger sequencing.

To clone IAV ORFs into the pCI mammalian expression vector, inserts were generated by PCR with primers that bound to the terminal regions of the PR8 NA/HA/NP ORFs and added *Eco*RI and *Sal*I restriction sites to the 5’/3’ ends respectively using the pDZ PR8 NA/HA/NP plasmids as templates. The PR8 PB1 ORF insert was generated with primers that bound to the terminal regions of the PR8 PB1 ORF and added *Mlu*I and *Kpn*I restriction sites to the 5’/3’ ends respectively using the pDZ PR8 PB1 plasmid as a template. Internal *Eco*RI/*Sal*I restriction sites in HA were removed via site-directed mutagenesis. The PR8 NA/HA/NP and PR8 PB1 inserts were then digested with the *Eco*RI/*Sal*I or *Mlu*I/*Kpn*I restriction enzymes respectively. The inserts were then ligated into the pCI vector, and transformed into chemically competent *E. coli* cells via the heat-shock method. Insert sequences were confirmed via sanger sequencing.

For epitope and eGFP-tagged NP expression vectors, mutations in PR8 NP were introduced via site directed mutagenesis. C-terminal 6x His tags were introduced by performing PCR with primers designed to add a C-terminal 6x His tag before the stop codon of the PR8 NP ORF using the pDZ PR8 NP plasmid as a template. A C-terminal HA-tag was added to PR8 PA by performing PCR with primers designed to add a C-terminal HA-tag before the stop codon of the PR8 PA ORF using the pDZ PR8 PA plasmid as a template. For cloning into the pCI plasmid, *Eco*RI/*Sal*I restriction sites were introduced to the 5’/3’ ends respectively of the PR8 NP-C-His (WT/F346S/R416A) and PR8 PA-HA tag ORFs. For cloning into the peGFP-C1 plasmid, *Bsp*EI/*Kpn*I restriction sites were introduced to the 5’/3’ ends respectively of the PR8 NP (WT/F346S/R416A) ORFs. The inserts were then digested with the *Eco*RI/*Sal*I (pCI PR8 NP:WT/F346S/R416A C-His and pCI PR8 PA-HA tag) or *Bsp*EI/*Kpn*I (peGFP-PR8 NP:WT/F346S/R416A) restriction enzymes, ligated into the pCI or peGFP vectors respectively, and transformed into chemically competent *E. coli* cells via the heat-shock method. Insert sequences were confirmed via sanger sequencing.

For lentiviral expression vectors, inserts were generated by PCR with primers designed to bind to the 5’ and 3’ terminal regions of the PR8 HA ORF and introduce *Bam*HI/*Not*I restriction sites to the 5’/3’ ends respectively using the pDZ PR8 HA plasmid as a template. The insert was then digested with the *Bam*HI/*Not*I restriction enzymes, ligated into the pHAGE-EF1aInt vector (generated by restriction digest of the pHAGE2-EF1aInt WSN HA W plasmid with the same restriction enzymes), and transformed into chemically competent *E. coli* cells via the heat-shock method. Insert sequences were confirmed via sanger sequencing.

### Cells

Madin-Darby canine kidney (MDCK), MDCK-SIAT1 cells, and 293T cells were obtained from Drs. Jonathan Yewdell, Jesse Bloom, and Joanna Shisler respectively and were maintained in Gibco’s minimal essential medium (MEM) with GlutaMax (Life Technologies) supplemented with 8.3% fetal bovine serum (Seradigm) (MEM+FBS) and incubated at 37°C with 5% CO_2_.

MDCK cells expressing the PR8 HA protein were generated via lentiviral transduction. 293T cells were transfected with 250ng each of the plasmids required for lentivirus generation (HDM Hgpm2, HDM tatlb, pRC CMV Rev1b, HDM VSV-G), and 1µg of the transfer vector pHAGE PR8 HA (generated as described above). One day post transfection, the transfection media was replaced with 2mL MEM+FBS. The next day, the lentiviral supernatant was collected and 1mL was used to infect three wells of MDCK cells plated in a 6 well plate at 10% confluency. Two days post transduction, the MDCK cells were harvested and combined, surface stained with an anti-HA antibody (H36-26 AF488), and positive cells were sorted out via fluorescence activated cell sorting (FACS).

### Viruses

Recombinant A/Puerto Rico/8/1934 (H1N1) (PR8) and A/Udorn/72 (H3N2) (Udorn) viruses were generated using 8 and 12 plasmid reverse genetics systems respectively. The rPR8 clones differ from the published sequence (GenBank accession no. AF389115 to AF389122) at two positions: PB1 A549C (K175N) and HA A651C (I207L) (numbering from initiating Met). Viruses containing single point mutations in the NP or NA segments were generated by rescuing the viruses using plasmids containing the specific mutations introduced via site-directed mutagenesis. The Udorn HA segment-encoding plasmids used were found to have the following mutations: A81G (N18D), C129T (H34Y), G1103T (silent), T1486A (F486Y), & A1614G (N529D) relative to the Udorn HA reference sequence (GenBank accession no. AX350190).

Viruses were rescued by transfecting 293T cells with 500ng each of the relevant reverse genetics plasmids using JetPrime (Polyplus) according to the manufacturer’s instructions. For the PR8 HA/NA chimeric, and PR8/Udorn NA chimeric PR8 NA ORF containing viruses with NP:F346S, the cells were also transfected with 500ng of the pCI PR8 NP, and the pCI PR8 HA/NA plasmids or pCI PR8 NA plasmid respectively to promote viral growth via expression of the native viral proteins. 18-24hrs post transfection, the media was replaced with viral growth media (MEM, 1 mM HEPES, 1 μg/mL TPCK trypsin (Worthington Biochemical Corporation; Lakewood, NJ, USA), 50 μg/mL gentamicin) containing 2×10^5^ MDCK cells). For the viruses with the chimeric PR8 HA/NA segments, the viral growth media was modified by adding 2×10^5^ PR8 HA+ MDCK cells instead of MDCK cells. Transfection supernatants were collected 24hrs post media change.

To generate the seed stocks of the PR8 NP:WT/F346S, Udorn NP:WT/F346S, PR8 Udorn HA/NA NP:WT/F346S, PR8 NP point mutant viruses, PR8 NA Codon Shuffle NP:WT/F346S viruses, PR8 NA:^C13U^ NP:WT/F346S, PR8 NA:^C4/13U^ NP:WT/F346S, PR8 NA:^Udorn UTR^ NP:WT/F346S, and PR8:Udorn HA,NA^PR8 ORF Proximal UTR^ NP:WT/F346S viruses, transfection supernatants were plaqued, and a single plaque was used to infect a single well of MDCK cells in a 6 well plate. Viral growth was performed in viral growth media (MEM, 1 mM HEPES, 1 μg/mL TPCK trypsin, 50 μg/mL gentamicin). Seed stocks were harvested and clarified (14000rpm, 15min, 4°C) between 24-72hrs post infection. Only seed stocks were generated for the PR8 NA Codon Shuffle NP:WT/F346S, PR8 NA:^C13U^ NP:WT/F346S, PR8 NA:^C4/13U^ NP:WT/F346S, PR8 NA:^Udorn UTR^ NP:WT/F346S, and PR8:Udorn HA,NA^PR8 ORF Proximal UTR^ NP:WT/F346S viruses. MDCK cells in a T75 or T175 flask were then infected with the seed stocks at an MOI of 0.001 or 0.01 TCID_50_/cell respectively, and the working stocks were harvested 24-72hrs post infection and clarified (3500rpm, 15min, 4°C). Viral growth was performed in viral growth media.

To generate the seed stocks of PR8 HA/NA chimeric viruses, PR8 HA+ MDCK cells in a single well of a 6 well plate were infected with 1mL of transfection supernatant for 1hr at 37°C with rocking, and then the transfection supernatant was removed and replaced with 3mL of viral growth media with 0.5 μg/mL TPCK trypsin and left to incubate for up to 48hrs. The infection supernatants were harvested and clarified (14,000rpm, 15min, 4°C), and 1mL of the infection supernatant was used to perform the next passage of the viruses in the PR8 HA+ MDCK cells. The passaging continued until cytopathic effect was observed. This occurred within the first two passages for all the viruses, and between 24-48hrs post infection. The PR8 HA/NA UTR swap NP:WT virus had a mutation in the PR8 HA ORF NA UTR segment (C33AàL5I).

To generate the seed stocks of the PR8 NA:^C4U^ NP:WT/F346S, PR8:NA^Udorn ORF proximal^ _UTR NP:WT/F346S, PR8 NA:C4U/Udorn ORF proximal UTR NP:WT/F346S, PR8 NA:C13U/Udorn ORF Proximal UTR NP:WT/F346S, PR8:NAUdorn 3’ ORF Proximal UTR NP:WT/F346S, PR8:NAUdorn 3’ UTR_ NP:WT/F346S, PR8 NA:^Udorn 5’ UTR^ NP:WT/F346S, and PR8:Udorn HA,NA^PR8 UTR^ NP:WT/F346S viruses, MDCK cells in a single well of a 6 well plate were infected with 1mL of transfection supernatant for 1hr at 37°C with rocking, and then the transfection supernatant was removed and replaced with 3mL of viral growth media and left to incubate for up to 48hrs. The infection supernatants were harvested and clarified (14,000rpm, 15min, 4°C) once CPE was observed.

Virus titers were determined by TCID_50_ assay on MDCK cells, or by determining the fraction of viral particles expressing NP (NPEU)(24) via flow cytometry on infected MDCK cells using the anti-NP AF647 (HB65) antibody.

### Next generation sequencing of viruses

Viral RNA was extracted from 140µL of the viral infection supernatant using the QIAamp Kit (Qiagen) and eluted in 60µL of Nuclease-free water (Ambion). Contaminating DNA was removed using the Qiagen RNase-free DNase Set, and then the RNA was cleaned using the RNeasy Kit (Qiagen) and eluted in 30µL of Nuclease-free water (Ambion). cDNA was synthesized using the Superscript III Reverse Transcriptase Kit (ThermoFisher) as follows: 1µL of 2µM MBTUni-12 primer (5’-ACGCGTGATCAGC<u>R</u>AAAGCAGG-3’) + 1µL 10mM dNTPs Mix (NEB #N0447S) + 8µL Nuclease-free water (Ambion) were added to 3µL of RNA and then incubated at 65°C for 5 min and then 4°C for 2 min. 4µL of 5x First Strand cDNA Synthesis Buffer + 1µL 0.1M DTT + 1µL SUPERase-In RNase Inhibitor (Invitrogen #AM2696) + 1µL Superscript III RT (Invitrogen #18080-044) were added to the reaction and then the reaction was incubated at 45°C for 50 min. The cDNA was then stored at −20°C. The PCR reaction to simultaneously amplify all eight gene segments was performed using Phusion polymerase (NEB #M0530L) as follows: 2.5µL of 10µM MBTUni-12 primer + 2.5µL of 10µM MBTUni-13 primer (5’-ACGCGTGATCAGTAGAAACAAGG-3’) + 10µL 5x HF Phusion Buffer + 1µL 10mM dNTPs mix (NEB #N0447S) + 0.5µL Phusion Polymerase + 28.5µL of Nuclease-free water (Ambion) was added to 5µL of cDNA. The cycling conditions for the PCR were as follows: 98°C for 30sec, (98°C for 10sec/ 57°C for 30sec/ 72°C for 1min 30sec) x 25, 72°C for 5min, 4°C Hold). The PCR products were cleaned using the Invitrogen PureLink PCR Purification Kit (Invitrogen #K310002) using the Buffer for the <300bp cutoff and eluted in 30µL of Nuclease-free water (Ambion). The PCR products were then subjected to next generation sequencing using the Illumina NovaSeq or MiSeq platforms.

### Time-course infections

MDCK cells were infected with the PR8 NP:WT/F346S or Udorn NP:WT/F346S viruses at an MOI of 0.1 NPEU/cell in a 24 well plate for 1hr at 37°C. 1hr post infection, the virus supernatant was replaced with 0.5mL MEM+FBS. 3hr post infection, the MEM+FBS was replaced with 0.5mL NH_4_Cl media (MEM, 50mM HEPES Buffer, 20mM NH_4_Cl, pH=7.2) to prevent viral spread. The cell monolayers were harvested 4, 8, and 12hrs post infection, and cellular RNA was extracted using the RNeasy kit (Qiagen).

Reverse transcription was performed using the Verso cDNA Synthesis Kit (ThermoFisher). The reactions were set up as follows: 4µL RNA + 4µL 5x cDNA Synthesis Buffer + 2µL dNTP Mix + 1µL 10µM PR8 RT_4A primer (5’-AGCAAAAGCAGG-3’) + 1µL RT Enhancer + 1µL Verso Enzyme Mix + 7µL Nuclease-free water, and incubated at 45°C for 50min, 95°C for 2min, and held at 4°C. cDNA was stored at −20°C. Quantitative real-time PCR on cDNA was carried out using Power SYBR green PCR Master Mix (Thermo Fisher) on a QuantStudio 3 thermal cycler (Thermo Fisher). The strand-specific forward and reverse primers for quantitative real-time PCR for PR8 HA, NP, and NA were 5’-AAGGCAAACCTACTGGTCCTGTT-3’ & 5’-AATTGTTCGCATGGTAGCCTATAC-3’, 5’-AGGCACCAAACGGTCTTACG-3’ & 5’-TTCCGACGGATGCTCTGATT-3’, and 5’-AAATCAGAAAATAACAACCATTGGA-3’ & 5’-ATTCCCTATTTGCAATATTAGGCT-3’ respectively. The strand-specific forward and reverse primers for Udorn HA, NP, and NA were 5’-GACTATCATTGCTTTGAGC-3’ & 5’-CACTAGTGTTCCGTTTGGC-3’ and 5’-CGGTCTTATGAACAGATGG-3’ & 5’-TCGTCCAATTCCATCAATC-3’ and 5’-AACAATTGGCTCTGTCTCTC-3’ & 5’-GTCGCACTCATATTGCTTG-3’ respectively. Reactions were set up as follows: 2µL cDNA + 10µL 2x Power SYBR Green MM + 1µL 10µM Forward Primer + 1µL 10µM Reverse Primer + 6µL Nuclease-free water. The cycling conditions were as follows: 50°C for 2min, 95°C for 10min, and then 95°C for 15 sec followed by 60°C for 1min repeated 40x.

### Analysis of primary viral transcription in infected cells

MDCK-SIAT1 cells were infected with the PR8 NP:WT/F346S viruses at an MOI of 5 TCID_50_/cell in a 6 well plate in the presence of 100µg/mL of cycloheximide (Sigma-Aldrich). Infected cells were harvested at 2 and 6hrs post-infection, and cellular RNA was extracted using the RNeasy Kit (Qiagen). Reverse transcription was performed using the Superscript III Reverse Transcriptase Kit (ThermoFisher). The reactions were set up as follows: 4µL RNA + 0.5µL 100µM Oligo dT_20_ primer (IDT) + 1µL 10mM dNTP + 6.5µL Nuclease-free water incubated at 65°C for 5min and then 4°C for 1min. 4µL 5x First Strand RNA Buffer + 1µL 0.1M DTT + 2µL SuperaseIN RNase Inhibitor (ThermoFisher) + 1µL of Superscript III Reverse Transcriptase was added to the previous reaction and incubated at 50°C for 60min, 70°C for 15min, and held at 4°C. cDNA was stored at −20°C. Quantitative real-time PCR on cDNA was carried out using Power SYBR green PCR Master Mix (Thermo Fisher) on a QuantStudio 3 thermal cycler (Thermo Fisher). The strand-specific forward and reverse primers for quantitative real-time PCR for PR8 HA and NA were as follows: PR8 HA 5’-AAGGCAAACCTACTGGTCCTGTT-3’ & 5’-AATTGTTCGCATGGTAGCCTATAC-3’ and PR8 NA 5’-AAATCAGAAAATAACAACCATTGGA-3’ & 5’-ATTCCCTATTTGCAATATTAGGCT-3’. Reactions were set up as follows: 1.5µL cDNA + 10µL 2x Power SYBR Green MM + 1µL 10µM Forward Primer + 1µL 10µM Reverse Primer + 6.5µL Nuclease-free water. The cycling conditions were as follows: 50°C for 2min, 95°C for 10min, and then 95°C for 15 sec followed by 60°C for 1min repeated 40x.

### 4SU RNA Pulse

60-70% confluent MDCK cells were infected with the PR8 NP:WT/F346S viruses at an MOI of 5 TCID_50_/cell and incubated at 37°C for 1hr. The virus was aspirated and then 3mL of MEM+FBS was added to each well. 7hpi the MEM+FBS was replaced with 1mL of fresh MEM+FBS containing 500µM 4-thiouridine (4SU) (Tri-Link Biotechnologies N-1025). The cells were kept in the dark during the labeling process to prevent cross-linking of 4SU to cellular proteins. 1hr post labeling the cells were harvested and cellular RNA was extracted using the RNeasy Kit (Qiagen).

The cellular RNA was then biotinylated by performing a reaction with EZ-Link HPDP-Biotin (ThermoScientific). Reactions conditions were as follows: 10µg RNA and Biotin-HPDP (0.2µg/µL final concentration) were added to Biotinylation Buffer (10mM Tris-HCl pH=7.5, 1mM EDTA) resulting in a total reaction volume of 250µL, and then incubated at room temperature with end-over-end rotation for 2hrs protected from light. RNA was then extracted using the chloroform: isoamyl alcohol procedure performed as follows: 400µL of chloroform: isoamyl alcohol (49:1 ratio) was added to each reaction, mixed, and then added to a 2mL Quanta Bio 5PRIME Phase Lock Heavy Tube. The phases were separated by centrifugation (Full speed, 5min, 4°C), and the aqueous layer (top) was transferred to a new 1.5mL microcentrifuge tube. The previous steps were then repeated one additional time. RNA was precipitated as follows: One volume of isopropanol, one-tenth volume of 5M NaCl, and 1µL of 15µg/mL GlycoBlue Coprecipitant (Invitrogen) were added to each sample and mixed. The samples were then frozen at −70°C overnight. The next day the samples were thawed and RNA was pelleted via centrifugation (Full speed, 20min, 4°C). The pellet was then washed 2x with 400µL of 80% ethanol (Full speed, 5min, 4°C). The pellet was then air-dried at room temperature for 5min and resuspended in 20µL of Nuclease-free water.

Biotinylated RNAs were then selectively purified using the µMACS Streptavidin Kit (Miltenyi Biotec) as follows: 15µL of RNA was added to 85µL of Nuclease-free water and denatured by incubating at 65°C for 10min followed by cooling on ice for 5min. 100µL of Miltenyi streptavidin beads were added to each reaction and incubated at room temperature for 15min with end-over-end rotation. Meanwhile, the µMACS columns were equilibrated by adding 100µL of the Equilibration Buffer for Nucleic Acid Applications and then washed with 1mL of wash buffer (100mM Tris-HCl pH=7.5, 10mM EDTA, 1M NaCl). The biotinylated RNA-streptavidin bead solution was then added to the columns, and then the columns were washed with 0.9mL of 65°C wash buffer 3x followed by 0.9mL of room temperature wash buffer 3x. The biotinylated RNA was then eluted with 150µL of 0.1M DTT. The RNA was stored at −70°C.

Reverse transcription was performed using the Superscript III Reverse Transcriptase Kit (ThermoFisher). A universal vRNA-specific primer or a tagged, segment-specific, vRNA-specific primer was used. The reactions were set up as follows: 2µL RNA + 0.5µL 10µM primer (Universal: PR8 RT_4A primer (5’-AGCAAAAGCAGG-3’) or segment specific: NA vRNA-24 tag (5’-GGCCGTCATGGTGGCGAATAATCCAAATCAGAAAATAACAACC-3’) or 10µM HA vRNA-36 tag primer (5’-GGCCGTCATGGTGGCGAATAAGGCAAACCTACTGGTCCTGTT-3’)) + 1µL 10mM dNTP + 8.5µL Nuclease-free water incubated at 65°C for 5min and then 4°C for 1min. 4µL 5x First Strand RNA Buffer + 1µL 0.1M DTT + 2µL SuperaseIN RNase Inhibitor (ThermoFisher) + 1µL of Superscript III Reverse Transcriptase were added to the previous reaction, and the reactions were incubated at 50°C for 60min, 70°C for 15min, and held at 4°C. cDNA was stored at −20°C. Quantitative real-time PCR on cDNA was carried out using Power SYBR green PCR Master Mix (Thermo Fisher) on a QuantStudio 3 thermal cycler (Thermo Fisher). The strand-specific forward and reverse primers for quantitative real-time PCR for PR8 HA and NA vRNA were: For RT reaction using universal primer (PR8 HA 5’-AAGGCAAACCTACTGGTCCTGTT-3’ & 5’-AATTGTTCGCATGGTAGCCTATAC-3’ and PR8 NA 5’-AAATCAGAAAATAACAACCATTGGA-3’ & 5’-ATTCCCTATTTGCAATATTAGGCT-3’), and for RT reaction using segment-specific primers (vtag (5’-GGCCGTCATGGTGGCGAAT-3’) & PR8 HA qPCR 3’ (5’-AATTGTTCGCATGGTAGCCTATAC-3’), and vtag (5’-GGCCGTCATGGTGGCGAAT-3’) & PR8 NA qPCR 3’ (5’-ATTCCCTATTTGCAATATTAGGCT-3’). Reactions were set up as follows: 1.5µL cDNA + 10µL 2x Power SYBR Green MM + 1µL 10µM Forward Primer + 1µL 10µM Reverse Primer + 6.5µL Nuclease-free water. The cycling conditions were as follows: 95°C for 10min, and then 95°C for 15 sec followed by 54/57°C for PR8 NA/HA vRNA respectively for 1min repeated 40x.

### Quantification of single replication cycle viral gene expression levels

MDCK cells were infected with viruses at an MOI of 0.1 NPEU/cell (0.03 NPEU/cell for PR8:Udorn HA,NA^PR8 UTR^ NP:WT/F346S or PR8:Udorn HA,NA^PR8 ORF Proximal UTR^ NP:WT/F346S viruses due to their low titer) or an MOI of 0.1 TCID_50_/cell (PR8 NP:WT v. F346S v. Y385A v. F479A) in a 24 well plate. 1hpi infection, the viral supernatant was removed and replaced with TCID_50_ media (MEM, 1 mM HEPES, 0.5 or 1 μg/mL TPCK trypsin, 50 μg/mL gentamicin). Cells were harvested at 8hpi and cellular RNA was extracted using the RNeasy Kit (Qiagen). For viruses derived from direct passage of the transfection supernatant in MDCK cells or a single infection with a plaque supernatant, the cellular RNA was treated with RNase-free DNaseI (Qiagen) and cleaned using the RNeasy Kit (Qiagen). Reverse transcription was performed using the Verso cDNA Synthesis Kit (ThermoFisher). The reactions were set up as follows: 4µL RNA + 4µL 5x cDNA Synthesis Buffer + 2µL dNTP Mix + 1µL 10µM PR8 RT_4A primer (5’-AGCAAAAGCAGG-3’) + 1µL RT Enhancer + 1µL Verso Enzyme Mix + 7µL Nuclease-free water, and incubated at 45°C for 50min, 95°C for 2min, and held at 4°C. cDNA was stored at −20°C. Quantitative real-time PCR on cDNA was carried out using Power SYBR green PCR Master Mix (Thermo Fisher) on a QuantStudio 3 thermal cycler (Thermo Fisher). The strand-specific forward and reverse primers for quantitative real-time PCR for PR8 HA, NP, and NA/PR8 NA ORF HA UTRs were 5’-AAGGCAAACCTACTGGTCCTGTT-3’ & 5’-AATTGTTCGCATGGTAGCCTATAC-3’, 5’-AGGCACCAAACGGTCTTACG-3’ & 5’-TTCCGACGGATGCTCTGATT-3’, and 5’-AAATCAGAAAATAACAACCATTGGA-3’ & 5’-ATTCCCTATTTGCAATATTAGGCT-3’ respectively. The strand-specific forward and reverse primers for Udorn HA, NP, and NA were 5’-GACTATCATTGCTTTGAGC-3’ & 5’-CACTAGTGTTCCGTTTGGC-3’ and 5’-CGGTCTTATGAACAGATGG-3’ & 5’-TCGTCCAATTCCATCAATC-3’ and 5’-AACAATTGGCTCTGTCTCTC-3’ & 5’-GTCGCACTCATATTGCTTG-3’ respectively.

The primers for the PR8 HA ORF NA UTR+Pack and PR8 NA ORF HA UTR+Pack segments were 5’-AAATCAGAAAATAACAACCATTGGA-3’ & 5’-CAACAATACCAACAGATTAGC-3’ and 5’-AAGGCAAACCTACTGGTCCTGTT-3’ & 5’-ATCAGCCCTACCACGAGGC-3’ respectively. The primers for the PR8 NA Codon Shuffle segment were: 5’-CAAATGGGACCGTCAAAGACCGC-3’ and 5’-GATGGGGCTTCACCGACTGG-3’. Reactions were set up as follows: 2µL cDNA + 10µL 2x Power SYBR Green MM + 1µL 10µM Forward Primer + 1µL 10µM Reverse Primer + 6µL Nuclease-free water. The cycling conditions were as follows: 50°C for 2min, 95°C for 10min, and then 95°C for 15 sec followed by 60°C for 1min repeated 40x.

### Flow cytometry to detect viral protein expression in singly-infected cells

MDCK cells were infected at an MOI of 0.03 TCID_50_/cell in a 6 well plate, or 1.5×10^6^ MDCK cells per well were infected with 10^-1^ to 10^-4^ dilutions of the viral stock in a 6 well plate (for NP-expressing unit (NPEU) determination). 1hpi the virus was replaced with 3mL of MEM+FBS. 3hpi the MEM+FBS was then replaced with 3mL of NH_4_Cl media (MEM, 50mM HEPES Buffer, 20mM NH_4_Cl, pH=7.2) to prevent secondary infection. 16hpi the cells were harvested, fixed and permeabilized with foxP3 fix/perm buffer (eBioscience). Cells were subsequently stained with one or multiple of the following antibodies: PR8 NA: Rabbit anti-NA (08-0096-03 EXSANG 3/9/09) followed by Donkey anti-Rabbit PE (711-116-152 Lot 121465), PR8 NP: HB65 AF647 or PacB, Udorn NA: Goat anti-N2 NA primary followed by Donkey anti-Goat (705-116-147) PE secondary, and Udorn HA: H14A2 AF647. The cells were run on a BD LSR II flow cytometer and analyzed using FlowJo version 10.1 (Tree Star, Inc.). Viral protein expression levels were determined from the geometric mean fluorescence intensity (GMFI) of the fluorophore associated with each protein. NPEU titers were calculated by dividing the number of infected cells (% NP+)(Total Number of Cells) by the dilution factor and the volume of the inoculum.

### PR8 NP:WT/F346S-C-His protein purification

100µg of the pCI PR8 NP:WT-C-His or pCI PR8 NP:F346S-C-His plasmids were transiently transfected into 100mL cultures of HEK Expi-293-F cells with ExpiFectamine according to the company protocol (Thermo Fisher). The transfected cells were pelleted (1000rpm, 5min, 4°C) and resuspended in 3mL of Equilibration Buffer (PBS, 10mM Imidazole, 1X cOmplete EDTA-free Protease Inhibitor Cocktail (Sigma-Aldrich)). Cells were kept frozen at −70°C until the next step of the purification procedure was performed. The cells were thawed and then lysed via the freeze-thaw method: Incubation in a dry-ice-ethanol bath followed by 42°C water bath repeated 2x. The chromosomal DNA was then sheared by passing the lysate through an 18G needle 4x. The cellular debris was pelleted by centrifugation (3000xg, 15min, 4°C), and the clarified lysates were transferred to new tubes. The clarified lysates were then treated with RNaseA (50µg/mL final concentration) for 2hrs at room temperature.

His-tagged proteins were then selectively purified using the HisPur Ni-NTA Spin Purification Kit, 0.2mL (ThermoFisher). To improve the purity of the eluted protein fractions and increase the protein yield, the eluate fractions for the PR8 NP:WT/F346S C-His proteins respectively were combined, the buffer for the eluate fractions was exchanged to Equilibration Buffer using Pierce Protein Concentrators (PES, 10kDa MWCO, 0.5mL) (ThermoFisher Scientific), and the HisPur Ni-NTA column purification was repeated once more. The eluate fractions for each protein were then combined, added to a Slide-A-Lyzer Dialysis Cassette (Extra Strength, 10kDa MWCO, 0.5-3mL capacity) (Thermo Scientific), and dialyzed in PBS at 4°C overnight. The dialyzed protein samples were concentrated using Pierce Concentrators (PES, 10kDa MWCO, 0.5mL) (Thermo Scientific), and the protein concentrations were determined using the Pierce Coomassie Plus Bradford Assay Kit (Thermo Scientific).

### Biolayer interferometry (BLI)

Binding kinetics of PR8 NP:WT/F346S C-His proteins to biotinylated single-stranded RNA (ssRNA) was determined by biolayer interferometry using an Octet Red96e instrument (FortéBio) at room temperature. Anti-streptavidin biosensors (FortéBio) were washed in 180µl 1X kinetics buffer (0.002% v/v Tween 20 in 1X PBS, pH 7.4) for 30min at room temperature. The experiment consisted of five steps: (1) baseline: 60s with 1X kinetics buffer; (2) loading: 300s with 5’ biotinylated 24nt ssRNA (5’-UUUGUUACACACACACACGCUGUG-3’) (IDT) at 1µM; (3) baseline: 60s with 1X kinetics buffer; (4) association: 60s with 50, 100, 200, and 400nM of PR8 NP:WT/F346S C-His protein; (5) dissociation: 300s with 1X kinetics buffer. Octet Data Acquisition (version 11.1, FortéBio) software was used to obtain biolayer interferometry data. To calculate the dissociation constant (KD) via curve fitting, a 1:1 binding model was used. Octet Data Analysis (version 11.1, FortéBio) software was used to analyze binding kinetics. Negative controls were set up with 1X kinetics buffer replacing 1µM of biotinylated ssRNA in the loading step.

### Transfection protocols for eGFP/His-tagged NP protein co-immunoprecipitation and HA-tagged vRNP complex pulldown experiments

For the eGFP/His-tagged NP protein co-immunoprecipitation experiment: 293T cells in a 10cm dish were transfected with 5µg each of one of the following pairs of plasmids using JetPrime (Polyplus) according to the manufacturer’s instructions: pCI PR8 NP:WT-C-His & peGFP-PR8 NP:WT, pCI PR8 NP:F346S-C-His & peGFP-PR8 NP:F346S, or pCI PR8 NP:R416A-C-His & peGFP-PR8 NP:R416A. For the HA-tagged vRNP complex pulldown experiment: 293T cells in a 10cm dish were transfected with 2µg each of the following plasmids (pCI PR8 PB2, pCI PR8 PB1, pCI PR8 PA-HA tag, pCI PR8 NP:WT/F346S/R416A, & pHH21 PR8 NA) using JetPrime (Polyplus) according to the manufacturer’s instructions.

### Co-immunoprecipitation

24hrs post transfection, the cells were lysed in MOPS Co-IP Lysis Buffer (20mM MOPS pH=7.5, 150mM NaCl, 0.5% Igepal CA-630,1x cOmplete EDTA-free Protease Inhibitor Cocktail (Sigma-Aldrich)) and clarified via centrifugation (20,000xg, 15min, 4°C). Mouse-anti-His (HIS.H8) antibody (Invitrogen) (for eGFP/His-tagged NP protein co-immunoprecipitation) or Mouse-anti-HA tag (2-2.2.14) antibody (Invitrogen) (for HA-tagged vRNP complex pulldown experiment) was added to the clarified lysates to a final dilution of 1:100, and the lysates were incubated with the antibody with end-over-end rotation overnight at 4°C. Antigen-antibody complexes were selectively purified using Pierce Protein A Agarose (ThermoFisher Scientific) according to the manufacturer’s instructions.

### Western blot

For eGFP/His-tagged NP protein co-immunoprecipitation experiment: Western blots were performed on the immunoprecipitated protein samples using the mouse-anti-eGFP (F56-6A.1.2.3) primary antibody (Invitrogen) (1:1,000) followed by the rat-anti-mouse-HRP conjugated (187.1) secondary antibody (BD Biosciences) (1:500) to detect co-immunoprecipitated PR8 NP:WT/F346S/R416A eGFP tagged proteins, or the mouse-anti-His (HIS.H8) primary antibody (Invitrogen) (1:1,000) followed by the rat-anti-mouse-HRP conjugated (187.1) secondary antibody (BD Biosciences) (1:500) to assess the pulldown efficiency of the PR8 NP:WT/F346S/R416A-C-His proteins. A western blot was also performed on the cell lysates using the rabbit-anti-NP primary antibody (GeneTex GTX125989) (1:1,000) followed by the goat-anti-rabbit-HRP conjugated (G-21234) secondary antibody (Invitrogen) (1:10,000) to detect the expression efficiencies of the his-tagged and eGFP-tagged PR8 NP proteins in transfected cells.

For HA-tagged vRNP-complex pulldown experiment: Western blots were performed on the immunoprecipitated protein samples using the mouse-anti-HA tag (2-2.2.14) primary antibody (Invitrogen) (1:1,000) followed by the rat-anti-mouse-HRP conjugated (187.1) secondary antibody (BD Biosciences) (1:500) to assess pulldown efficiency of the HA-tagged PA, or the rabbit anti-NP polyclonal antibody (GeneTex GTX125989) (1:1,000) followed by the goat-anti-rabbit-HRP conjugated (G-21234) secondary antibody (Invitrogen) (1:10,000) to detect co-immunoprecipitated NP.

Proteins were visualized using the SuperSignal Pico West Plus Chemiluminescent Substrate (ThermoFisher Scientific) and imaged using the iBright CL1000 Imaging System (Invitrogen).

### Quantifying gene segment ratios in viral stocks

140µL of the viral stock was treated with 0.25µg of RNaseA for 30min at 37°C. The viral RNA was then extracted using the QIAamp Viral RNA Extraction Kit (Qiagen) and eluted in 60µL of nuclease-free water (Ambion). RNA was treated with RNase-free DNaseI (Qiagen) and cleaned using the RNeasy Kit (Qiagen). Reverse transcription was performed using the Verso cDNA Synthesis Kit (ThermoFisher). The reactions were set up as follows: 4µL RNA + 4µL 5x cDNA Synthesis Buffer + 2µL dNTP Mix + 1µL 10µM PR8 RT_4A primer (5’-AGCAAAAGCAGG-3’) + 1µL RT Enhancer + 1µL Verso Enzyme Mix + 7µL Nuclease-free water, and incubated at 45°C for 50min, 95°C for 2min, and held at 4°C. cDNA was stored at −20°C. Quantitative real-time PCR on cDNA was carried out using Power SYBR green PCR Master Mix (Thermo Fisher) on a QuantStudio 3 thermal cycler (Thermo Fisher). The strand-specific forward and reverse primers for quantitative real-time PCR for PR8 HA, NP, and NA/PR8 NA ORF HA UTRs were 5’-AAGGCAAACCTACTGGTCCTGTT-3’ & 5’-AATTGTTCGCATGGTAGCCTATAC-3’, 5’-AGGCACCAAACGGTCTTACG-3’ & 5’-TTCCGACGGATGCTCTGATT-3’, and 5’-AAATCAGAAAATAACAACCATTGGA-3’ & 5’-ATTCCCTATTTGCAATATTAGGCT-3’ respectively. The strand-specific forward and reverse primers for Udorn HA, NP, and NA were 5’-GACTATCATTGCTTTGAGC-3’ & 5’-CACTAGTGTTCCGTTTGGC-3’ and 5’-CGGTCTTATGAACAGATGG-3’ & 5’-TCGTCCAATTCCATCAATC-3’ and 5’-AACAATTGGCTCTGTCTCTC-3’ & 5’-GTCGCACTCATATTGCTTG-3’ respectively. The primers for the PR8 HA ORF NA UTR+Pack and PR8 NA ORF HA UTR+Pack segments were 5’-AAATCAGAAAATAACAACCATTGGA-3’ & 5’-CAACAATACCAACAGATTAGC-3’ and 5’-AAGGCAAACCTACTGGTCCTGTT-3’ & 5’-ATCAGCCCTACCACGAGGC-3’ respectively. The primers for the PR8 HA ORF NA UTR segment were 5’-CAGGAGTGCCAAATTGAGGATGG-3’ and 5’-CCGGCAATGGCTCCAAATAGACC-3’. Reactions were set up as follows: 2µL cDNA + 10µL 2x Power SYBR Green MM + 1µL 10µM Forward Primer + 1µL 10µM Reverse Primer + 6µL Nuclease-free water. The cycling conditions were as follows: 50°C for 2min, 95°C for 10min, and then 95°C for 15 sec followed by 60°C for 1min repeated 40x.

## Acknowledgments

This research was funded in whole, or in part, by the Wellcome Trust [FC011104]. For the purpose of Open Access, the author has applied a CC BY public copyright license to any Author Accepted Manuscript version arising from this submission. We would like to thank Tongyu Liu and Jiayi Sun for providing plasmids and members of the Brooke lab for their feedback on this manuscript. Additionally, we would like to thank Sonya Kumar Bharathkar for assistance with mammalian protein expression in HEK Expi-293-F cells and Sarah Leonard for helpful discussions regarding NP-RNA binding experiments. We are also grateful to Drs. Ervin Fodor and Andy Mehle for helpful discussions. This work has been generously supported by the National Institute of Allergy and Infectious Diseases of the National Institutes of Health under awards K22AI116588 and R01AI139246, the Roy J. Carver Charitable Trust under award 17-4905, the Francis Crick Institute which receives its core funding from Cancer Research UK (FC011104), the UK Medical Research Council (FC011104), and the Wellcome Trust (FC011104), and startup funds from the University of Illinois.

## SI figure legends

***S1 Fig. Quantifying the abundance of newly synthesized, 4SU-labeled vRNAs using vRNA and segment-specific primers during the cDNA synthesis and qPCR steps.** Normalized abundance of 4SU-labeled NA vRNA in MDCK cells infected with PR8 NP:WT/F346S at an MOI of 5 TCID_50_/cell for 7hrs and pulsed with 500µM of 4SU for 1hr as determined by RT-qPCR using a tagged, vRNA and segment-specific primer during the cDNA synthesis step, and a primer pair consisting of a tag-specific primer and segment-specific primer for the qPCR step. Each data point represents a single cell culture well replicate from a single experiment.*

***S2 Fig. Quantification of gene segment ratios in viral RNA stocks.** 140µL of viral RNA supernatant was treated with 0.25µg RNaseA, viral RNA was extracted and DNase-treated, and then viral gene segment abundance was quantified using RT-qPCR. A/B.) Normalized HA ORF (A) or NA ORF (B) containing viral RNA abundance in the viral stocks of the PR8 HA/NA UTR+Pack Swap NP:WT/F346S and PR8 HA/NA UTR Swap NP:WT/F346S viruses. C.) Normalized viral RNA abundance in the viral stocks of the Udorn NP:F346S, PR8:Udorn HA, NA NP:F346S, PR8:Udorn HA, NA^PR8 UTR^ NP:F346S, and PR8:Udorn HA, NA^PR8 ORF Proximal UTR^ NP:F346S viruses relative to Udorn NP:WT, PR8:Udorn HA, NA NP:WT, PR8:Udorn HA, NA:^PR8 UTR^ NP:WT, and PR8:Udorn HA, NA^PR8 ORF Proximal UTR^ NP:WT viruses respectively. D.) Normalized viral RNA abundance in the viral stocks of the indicated viruses with NP:F346S as determined by RT-qPCR normalized to the NP:WT versions of each virus. Each data point represents a qPCR technical replicate.*

