## Supplemental Figures for "Interactions between influenza A virus nucleoprotein and gene segment UTRs facilitate selective modulation of viral gene expression"

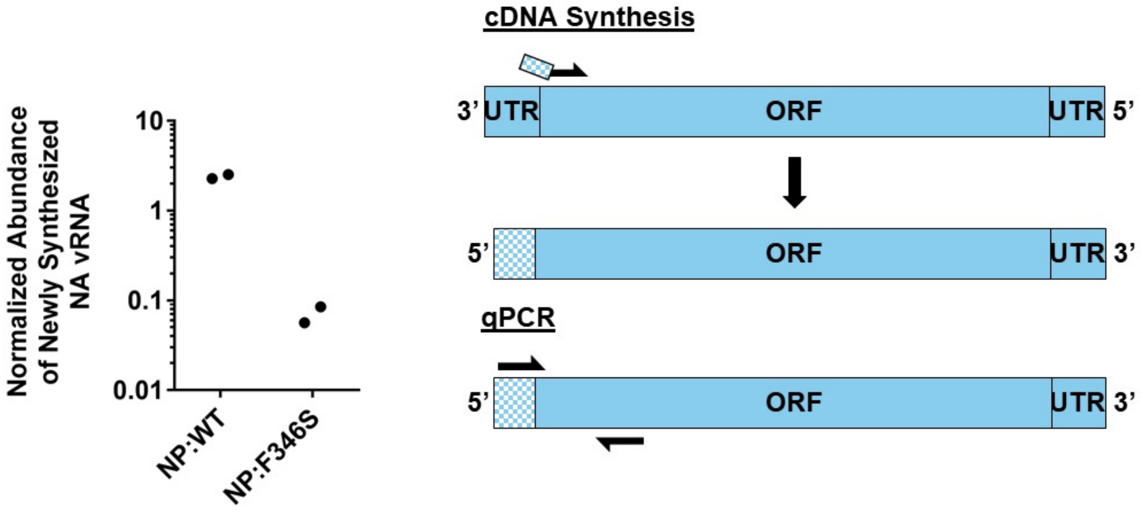


***S1 Fig. Quantifying the abundance of newly synthesized, 4SU-labeled vRNAs using vRNA and segment-specific primers during the cDNA synthesis and qPCR steps.***
*Normalized abundance of 4SU-labeled NA vRNA in MDCK cells infected with PR8 NP:WT/F346S at an MOI of 5 $TCID_{50}$/cell for 7hrs and pulsed with 500µM of 4SU for 1hr as determined by RT-qPCR using a tagged, vRNA and segment-specific primer during the cDNA synthesis step, and a primer pair consisting of a tag-specific primer and segment-specific primer for the qPCR step. Each data point represents a single cell culture well replicate from a single experiment.*

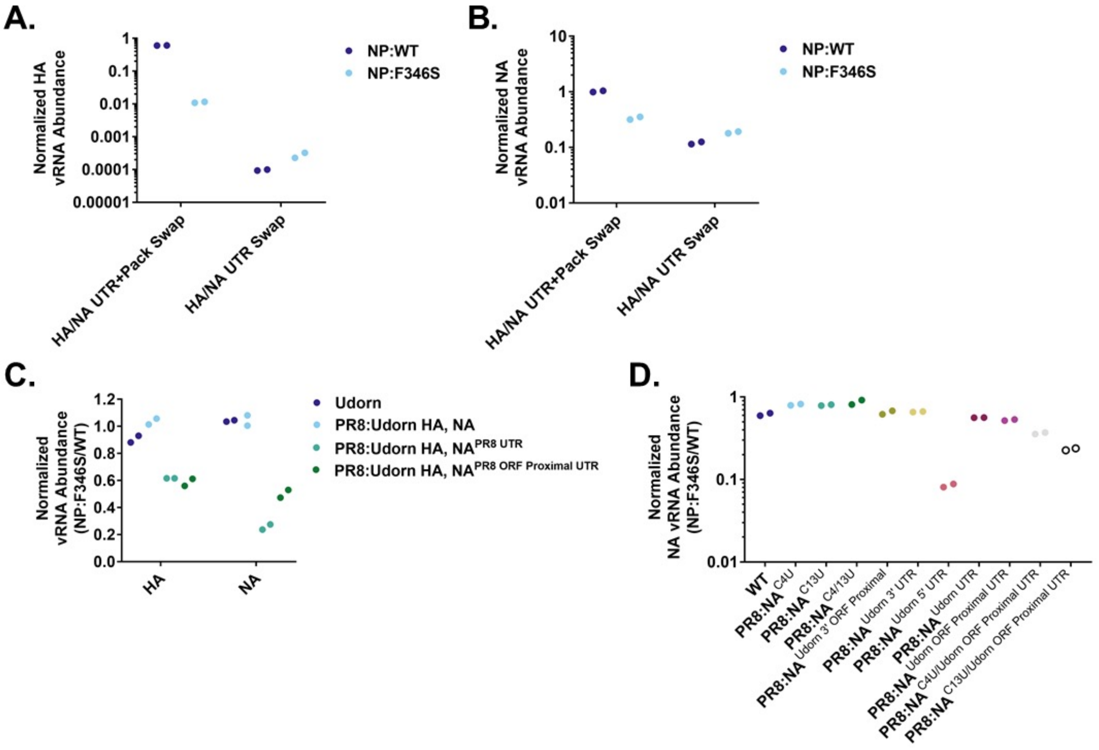


***S2 Fig. Quantification of gene segment ratios in viral RNA stocks.*** *140µL of viral RNA supernatant was treated with 0.25µg RNaseA, viral RNA was extracted and DNase-treated, and then viral gene segment abundance was quantified using RT-qPCR.* ***A/B.)*** *Normalized HA ORF* ***(A)*** *or NA ORF* ***(B)*** *containing viral RNA abundance in the viral stocks of the PR8 HA/NA UTR+Pack Swap NP:WT/F346S and PR8 HA/NA UTR Swap NP:WT/F346S viruses.* ***C.)*** *Normalized viral RNA abundance in the viral stocks of the Udorn NP:F346S, PR8:Udorn HA, NA NP:F346S, PR8:Udorn HA, NA*$^{PR8\ UTR}$ *NP:F346S, and PR8:Udorn HA, NA*$^{PR8\ ORF\ Proximal\ UTR}$ *NP:F346S viruses relative to Udorn NP:WT, PR8:Udorn HA, NA NP:WT, PR8:Udorn HA, NA:*$^{PR8\ UTR}$ *NP:WT, and PR8:Udorn HA, NA*$^{PR8\ ORF\ Proximal\ UTR}$ *NP:WT viruses respectively.* ***D.)*** *Normalized viral RNA abundance in the viral stocks of the indicated viruses with NP:F346S as determined by RT-qPCR normalized to the NP:WT versions of each virus. Each data point represents a qPCR technical replicate.*
